# Naloxone as mitochondrial phenotype rescuer in a 3D bioprinted LCHADD/VLCADD model

**DOI:** 10.64898/2026.09.07.749837

**Authors:** Antonia Degen, Michael Ausserlechner, Verena Sturmlehner, Christian Ploner, Sara Tucci, Daniela Karall, Hamed Homaeirad, Lukas Neumann, Judith Hagenbuchner

## Abstract

For patients diagnosed with long-chain 3-hydroxyacyl-CoA dehydrogenase deficiency (LCHADD) or very-long-chain acyl-CoA dehydrogenase deficiency (VLCADD), fasting episodes or high-energy demands remain life threatening. Due to the low incidence, clinical trials for novel LCHADD/VLCADD therapies are limited, and current mouse models recapitulate human symptoms only partially. Here, we report the use of mitochondrial morphology and 3D bioprinted, vascularized tissue models to establish a robust testing platform for dietary-based and experimental treatment approaches. Using this platform, we demonstrated that mitochondrial morphology is strictly regulated by NOX2-driven ROS formation. Treatment of LCHADD/VLCADD-derived fibroblasts with the NOX2-inhibitor naloxone led to the reassembly of mitochondrial structures controlled by DNM1L/MFN2. Using RNA transcriptomics, we identified a pro-fibrotic phenotype in LCHADD and VLCADD patient cells, which impaired vessel formation in fully 3D bioprinted human tissue equivalents. Metabolic supplementation with dietary approaches, which are used in standard therapy, partially improved vascularization. Naloxone induced the strongest improvement, restoring vessel length and network complexity to those of healthy controls, suggesting increased oxidative stress as main driver. Interestingly, naloxone had no effect on healthy fibroblasts, underscoring its safety. Taken together, these findings suggest the opioid antagonist naloxone as a potential rescue medication during LCHADD/VLCADD-driven metabolic crises.

## Introduction

Long-chain 3-hydroxyacyl-CoA deficiency (LCHADD) and very-long-chain acyl-CoA dehydrogenase deficiency (VLCADD) are rare inherited metabolic disorders of long-chain fatty acid oxidation (LC-FAOD), with an estimated incidence of approximately 1 in 50,000 to 1 in 100,000. Both disorders present with clinically severe, potentially life-threatening manifestations, which occur particularly during metabolic stress such as fasting or illness. Mainly affected are high-energy demand tissues including heart and skeletal muscle [1–5]. Implementing the newborn screening has improved early diagnosis. However, morbidity and mortality remain significant [6].

Current treatment strategies are based on restricting long-chain fatty acids in the diet, supplementing medium-chain triglycerides, and avoiding catabolic states. In selected patient cohorts, anaplerotic therapy using an odd-carbon medium-chain triglyceride triheptanoin (C7) has shown clinical benefits [7, 8]. These interventions aim to bypass the metabolic block in long-chain fatty acid oxidation and to replenish tricarboxylic acid cycle intermediates, maintaining cellular energy metabolism [9].

Despite these therapeutic advances, disease mechanisms are still not fully understood. The low incidence of LCHADD and VLCADD limits the availability of sufficiently large patient cohorts for clinical studies. A Recently developed LCHADD knock-in mouse exhibit more severe symptoms and a phenotype more similar to the situation in humans than former mouse models, which might also contribute to further understanding of LCHADD pathogenesis [10]. There is increasing evidence that mitochondrial dysfunction and oxidative stress substantially contribute to disease pathophysiology beyond the primary defect in fatty acid oxidation. Since mitochondria regulate not only cellular energy metabolism, but also redox homeostasis, cytoskeletal organization, and stress-responsive signaling pathways, persistent mitochondrial dysfunction may have consequences that extend beyond bioenergetic impairment [11–13].

Before, we demonstrated that patient-derived fibroblasts from individuals with LCHADD and VLCADD exhibit altered mitochondrial network organization, characterized by increased fragmentation and reduced connectivity [11]. These changes are associated with impaired mitochondrial dynamics, reduced oxidative phosphorylation, increased reliance on glycolysis, and elevated levels of intracellular reactive oxygen species (ROS). NADPH oxidase 2 (NOX2) has been mechanistically implicated in ROS accumulation, as its inhibition reduces oxidative stress and restores mitochondrial morphology [11]. In line with these findings, mitochondria-targeted antioxidants partially restored redox balance and bioenergetic function in patient-derived VLCADD fibroblasts [14]. Together, these findings suggest that ROS represents a driver of disease-associated cellular dysfunction and provides rationale for therapeutically targeting redox imbalance.

On this basis, pharmacological modulation of NOX2-dependent ROS represents a potential therapeutic strategy. Besides its established clinical use as an opioid receptor antagonist, naloxone has been reported to inhibit NOX2-dependent ROS production independent of opioid receptor signaling [15, 16].

In this study, we investigated whether metabolic and redox-targeted interventions can reverse the disease-associated phenotype of LCHADD and VLCADD fibroblasts. Specifically, we compared established metabolic interventions with pharmacological inhibition of NOX2-dependent ROS production and examined their effects on mitochondrial organization, cellular metabolism, transcriptional remodeling, cytoskeletal organization, and fibroblast function. To determine whether these cellular changes translate into tissue level, we developed a fully 3D bioprinted vascularized tissue model combining patient-derived fibroblasts with endothelial cells and adipose-derived stromal cells. This approach enabled us to evaluate how restored fibroblast homeostasis influences extracellular matrix remodeling and vascular network formation within a human tissue-like microenvironment.

## Materials and Methods

### Primary cells and used cell lines

Primary skin fibroblasts were obtained from a healthy pediatric donor (male) and from patients diagnosed with LCHADD (n = 3; 2 male and 1 female) or VLCADD (n = 1; female) at the metabolic centers in Vienna, Graz and Innsbruck as previously described [7] [11]. Fibroblasts were cultured in high-glucose Dulbecco’s Modified Eagle Medium (DMEM) supplemented with 10% fetal bovine serum (FBS; Thermo Fisher Scientific, USA), 100 U/ml penicillin, 100 µg/ml streptomycin, 2 mM L-glutamine. Primary human adipose-derived stem cells (ASCs) were isolated from abdominal subcutaneous adipose tissue obtained from patients undergoing elective abdominoplasty at the Department of Plastic and Reconstructive Surgery, Medical University of Innsbruck [17, 18]. The study was approved by the Ethics Committee of the Medical University of Innsbruck (EK0244/2018), and written informed consent was obtained from all participating donors. ASCs were cultured as described by Ploner et al [19]. Human dermal microvascular endothelial cells (HDMVECs; ATCC® PCS-110-010™) and human umbilical vein endothelial cells (HUVECs; ATCC® CRL-1730™) were cultured in F-12K Nutrient Mixture (Thermo Fisher Scientific, Waltham, USA) supplemented with 10% FBS, 100 U/ml penicillin, 100 µg/ml streptomycin, 30 µg/ml endothelial cell growth supplement (ECGS; Thermo Fisher Scientific, Waltham, USA), and 100 µg/ml heparin. HUVECs and HDMVECs were supplemented with 2 mM or 4 mM L-glutamine, respectively. All cells were maintained at 37 °C in a humidified incubator with 5% CO2 and 95% relative humidity and routinely tested for mycoplasma contamination (Minerva Biolabs, Berlin, Germany).

### Expression vectors and genetically modified cell lines

Retroviral and lentiviral particles were produced and used for the genetically modification of endothelial cells and patient-derived fibroblasts, respectively. The cell lines HDMVEC/hTert-EYFP and HUVEC/hTert-EYFP have been described by Nothdurfter et al [20]. Lentiviral transduction of fibroblasts was performed as previously described [21] using pLKO-GFPmt, a GFP-tagged Cox8 plasmid provided by T. Miki [22], to enable visualization of mitochondrial structures.

### Reagents

The following reagents were used for treatment experiments: C7 (triheptanoin/UX007; Ultragenyx Pharmaceutical, Novato, USA), linoleic acid, glucose, and naloxone (Amomed, Vienna, Austria). Fatty acid-BSA complexes for supplementation with C7 and linoleic acid were prepared as previously described [23]. Unless otherwise specified, all reagents were obtained from Merck (Darmstadt, Germany).

### Glucose determination

Glucose consumption was quantified after cultivation of equal cell numbers for 72 hours using a colorimetric assay (Glucose (GO) Assay Kit) according to the manufacturer’s instructions. Absorbance was measured at 450 nm using a Spark® Cyto cell multimode plate reader (Tecan, Männedorf, Switzerland). Glucose levels were normalized to the cell number determined at the experimental endpoint. Data were collected from three independent experiments per condition.

### Live cell fluorescence microscopy and immunofluorescence

All images were acquired using a Zeiss Axiovert200M microscope equipped with an ApoTome.2 system. Cellular fluorescence intensity was quantified within 30x30 µm regions using Axiovision software (Zeiss, Vienna, Austria). For ROS measurement, cells were cultured on collagen-coated (rat-tail collagen I, 0.1 mg/ml; BD Bioscience, San Jose, USA) 8-well µ-Slides (ibidi GmbH, Gräfelfing, Germany) and incubated with CM-H2XROS (500 nM; Invitrogen, Carlsbad, USA) as previously described [11]. For live-cell analysis of GFP-tagged mitochondrial structures, fibroblasts were cultured on collagen-coated glass slides for five days. For quantitative analysis, 30x30 µm regions of at least 60 single mitochondria were analyzed for mitochondrial branches and dots automatically using our recently developed MiMoQuant software (currently in revision for Bioinformatics Advances). For immunofluorescence analysis, cells were cultured on collagen-coated 8-well µ-Slides. Prior to fixation with 4% paraformaldehyde, cells were stained with CMXRos (300 nM; Invitrogen, Carlsbad, USA). Cells were permeabilized with 0.1% Triton-X 100/PBS and blocked with 1% FBS, 2% BSA in PBS to prevent unspecific binding. Primary antibody incubation was performed overnight at 4 °C using antibodies against dynamin-related protein 1 (DNM1L; Abcam, Cambridge, UK) and mitofusin 2 (MFN2; Invitrogen, Carlsbad, USA). Primary antibodies were detected using goat anti-rabbit and goat anti-mouse secondary antibodies (Invitrogen, Carlsbad, USA). Nuclei were counterstained with Hoechst-33342 (10 µg/ml). Each dataset included at least 10 individual cells per condition. For cytoskeletal staining, cells were fixed and permeabilized as described above for immunofluorescence analysis. F-actin was stained with Phalloidin-Atto590 (50 nM, Thermo Fisher Scientific, Waltham, USA), and nuclei were counterstained with Hoechst-33342. Three independent regions were analyzed per condition.

### Subcellular fractionation and immunoblotting

Cytoplasmic and heavy membrane extracts were prepared as previously described [21, 24] and separated by SDS-PAGE (10 µg/lane) and Western blotting. Membranes were blocked, incubated with primary antibodies against α-Tubulin and CoxIV as separation controls (Cell Signaling Technology Inc., Boston, USA), as well as OXPHOS, DNM1L, and MFN2 (Abcam, Cambridge, UK), washed and detected with secondary antibodies. The blots were developed either by enhanced chemiluminescence (Cytiva, Marlborough, USA) or by Alexa488- and Alexa568-labeled secondary antibodies and imaged with an iBright imaging system (Thermo Fisher Scientific, Waltham, USA). iBright™ analysis software (Thermo Fisher Scientific, Waltham, USA) was used for gel quantification according to the user’s manual.

### RNA isolation

Total RNA was isolated from equal cell numbers following 72 h of culture with or without naloxone treatment using the Quick-RNA Miniprep Plus Kit (Zymo Research, Irvine, USA) according to the manufacturer’s instructions.

### RNA sequencing and gene expression analysis

Library preparation, sequencing and gene expression analysis were performed at the company XPseq Analytics GmbH (Austria) using their ExpressoSeq rapid 3’ RNA-Seq service. Sequencing was performed on a PromethION 2 Solo sequencer on a R10 flow cell and the Ligation Sequencing Kit V14 (Oxford Nanopore Technologies). Sequencing reads were aligned to the human (GENCODE Release 47, GRCh38.p14) reference genome using the Minimap2 aligner (version 2.28). Read counting and differential gene expression analysis was performed with the Rsubread (version 2.22.1) and edgeR (version 4.6.3) packages in R (version 4.5.0). To account for baseline differences across the unique patient-derived cell lines, differential expression was modeled using a blocked generalized linear model with cell line identity as a blocking factor. Gene set enrichment analysis was performed using the Camera function with the MSigDB Hallmark gene sets (version v2024.1.Hs). Data visualization was performed in R (version 4.6.0) using RStudio (version 2026.04.0.526). Volcano plots were generated with the EnhancedVolcano package (version 1.30.0), while heatmaps were generated using the pheatmap package (version 1.0.13). Ccolor gradients were implemented using the color space package (version 2.1-2).

### Hydrogel preparation and rheometry

Gelatin-methacryloyl (GelMA) was produced as recently described by Shirahama et al [25]. The viscoelastic properties of the hydrogels were measured using ElastoSense™ Bio (Rheolution, Montreal, Canada). For rheometric analysis, 3 ml hydrogel consisting of 4%, 4.5%, or 5% gelatin-methacrylate (GelMA) (w/v) and 0.25% LAP (w/v) were prepared and analyzed in “stiff mode” to determine the shear storage modulus (G’). Bioinks were gelled in the rheometer by exposure to 405 nm violet light. Each dataset included at least three independent measurements per concentration. The degree of GelMA functionalization was routinely assessed for each production batch by TNBS assay (0.1% (w/v) TNBS; 2,4,6-trinitrobenzene sulfonic acid) as described by Zhu et al [26]. The average degree of functionalization of hydrogels prepared for rheological measurements was 96.6%.

### Biofabrication of 3D LCHADD/VLCADD models

The 3D bioprinted models composed of healthy or patient-derived fibroblasts, endothelial cells, and stem cells were biofabricated using a custom-prepared, standardized functionalized GelMA hydrogel. The specialized hydrogel formulation optimized for vascularized tissues were adapted from Nothdurfter et al [20, 25]. Prior to bioprinting, a ring-shaped support structure was produced by direct dispensing of the thermosensitive material Pluronic F-127 (35% w/v) [27]. Printing was conducted at room temperature with a 20 G, 1” needle (Gonano Dosiertechnik, Kirchdorf a.d.K., Austria), with an applied pressure of 160 kPa and a feed rate of 4 mm/s. The resulting Pluronic rings had a diameter of 0.58 mm and a height of 0.4 mm. All printing processes were performed using a multi-printhead 3D Discovery bioprinter (CF300H, 3D Discovery BioSafety, RegenHu, Villmergen, Switzerland). Cell-laden hydrogels were subsequently printed into the preformed Pluronic rings using a micro-jet printhead operated at 10 kPa and a feed-rate of 15 mm/s. An infill grid spacing of 1 mm and a layer height of 0.05 mm were applied. The hydrogel consisted of 4.0 % GelMa and 0.25 % LAP and contained 12.5x10^6^ cells/ml. Cells were mixed at a ratio of 4:3:1:2, comprising ASCs, HDMVEC/hTert-EYFP, HUVEC/hTert-EYFP, and either healthy or patient-derived fibroblasts. Each printed construct was photocrosslinked immediately after jetting of each layer for 5 s using 390-400 nm light at 360 mW. Upon completion of the printing process, a final crosslinking step was performed for 30 seconds at 405 nm (50 mW/cm^2^). Subsequent cross-linking of the hydrogel components and cultivation of the vascularized tissue models were carried out as previously described [20].

### Characterization and quantification of vessel formation

Spontaneous vessel formation in 3D bioprinted tissue surrogates was analyzed by fluorescence microscopy. Images were acquired using a Zeiss Axiovert200M microscope (Zeiss, Vienna, Austria). The dataset included at least three independent experiments per treatment, with each experiment comprising two individual tissue surrogates. Vessel formation was quantified using AngioTool software [28]. Image acquisition and threshold calibration were standardized using untreated healthy control constructs and applied uniformly across all conditions. The software extracted vessel skeletons and branching points, enabling reproducible comparison of network complexity, branching density, and overall vascular organization between experimental groups. A minimum of two microscopic images were analyzed for each tissue surrogate.

### Statistics

Data analysis was performed using unpaired Student’s t–test, one-way and two-way ANOVA test combined with Šídák’s test or Dunnett’s multiple test for multiple comparisons. Data are presented as mean ± standard error of the mean (SEM). A p-value less than 0.05 was considered statistically significant, with levels of significance indicated as follows: *p* < 0.05 (*), *p* < 0.01 (**), *p* < 0.001 (***), and *p* < 0.0001 (****). Prism11 (GraphPad PRISM Software Inc., version 11.0.0) was used for statistical analysis.

## Results

### Rescue of mitochondrial network structure and metabolic function in LCHADD and VLCADD patient-derived fibroblasts

In a previous study, we already identified ROS-dependent mitochondrial structural abnormalities in patient-derived LCHADD and VLCADD fibroblasts which could be restored by the NOX inhibitor Phox-I [11]. Based on these observations, we analyzed mitochondrial structures under different metabolic or redox-modulating treatments using fibroblasts isolated from skin biopsies of LCHADD and VLCADD patients [7] infected with a COX8-targeted GFP reporter. Under baseline conditions, patient-derived fibroblasts consistently displayed a fragmented, punctate mitochondrial morphology as previously described for LCHADD and VLCADD cells [11], in contrast to the elongated and highly interconnected networks observed in fibroblasts from healthy donor fibroblasts (Figure 1A). These findings confirm that mitochondrial network disruption represents a consistent and reliable cellular hallmark of LCHADD and VLCADD fibroblasts. Cells were exposed to candidate compounds for three days, which were selected based on their clinical relevance or mechanistic basis. These included: C7 (triheptanoin, 2.5 mmol/L), a synthetic medium-chain triglyceride used in the clinical management of LC-FAOD, including LCHADD and VLCADD [7, 11, 28]; linoleic acid (LA; 25 µM), included in LC-FAOD nutrition guidelines to prevent essential fatty acid deficiency during a restricted long-chain fatty acid diet [29]; high-glucose (9 g/L) supplementation, to address altered substrate utilization [11]; and the NOX2 inhibitor naloxone (1 µM, every 12 h), targeting elevated ROS production previously linked to NOX2 activity in patient cells [11]. Remarkably, all treatment conditions induced the reassembly of the mitochondrial network, resulting in a visible shift toward elongated, interconnected structures (Figure 1A). We used a recently developed annotation tool, MiMoQuant (currently in revision for Bioinformatics Advances), which incorporates YoloX-based object detection, to quantify branches and isolated mitochondria. Supplemental Figure 1 illustrates how the tool identifies branches (green) and isolated mitochondria (red) within a selected ROI in LCHADD-derived fibroblasts. All treatment conditions were associated with an increase in the number of mitochondrial branches and a reduction in the number of mitochondrial fragments, reflecting increased network connectivity (Figure 1B). These data demonstrate that the mitochondrial structural defects observed in LCHADD and VLCADD fibroblasts are reversible and can be modulated pharmacologically.

**Figure 1.**
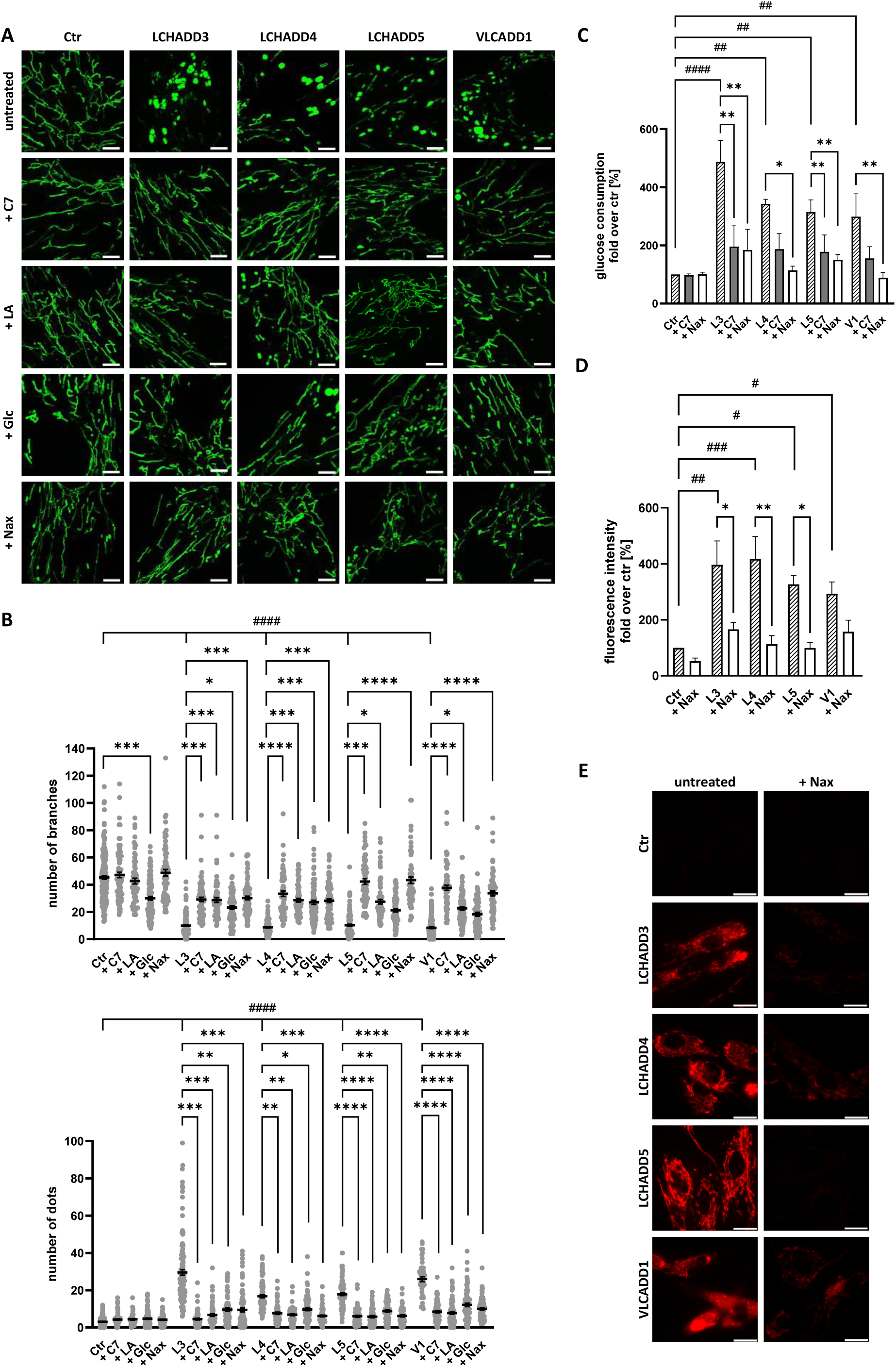
Rescue of mitochondrial network structure and metabolic function in LCHADD and VLCADD patient-derived fibroblasts. **A** Representative live-cell fluorescence images of mitoGFP-transfected healthy donor (ctr) or patient-derived fibroblasts with or without C7 (2.5 mmol/L), linoleic acid (LA; 25 µM), glucose (Glc; 4.5 g/L), or naloxone (Nax; 1 µM, every 12 h) (72 h). Scale bars, 5 µm. **B** Quantification of mitochondrial branches and dots in 30 × 30 µm regions of interest (≥ 60 mitochondria from 3-4 independent experiments per condition) under the conditions shown in A. **C** Glucose consumption in untreated, C7- or naloxone-treated fibroblasts (72 h). Data are expressed as fold change relative to untreated control cells (n = 4). **D** Quantification of ROS-associated fluorescence intensity under the conditions shown in **E**, expressed as fold change relative to untreated control fibroblasts (n = 3-4 independent experiments per condition). **E** Representative fluorescence images of intracellular ROS detected with CM-H₂XROS in untreated or naloxone-treated fibroblasts (72 h). Images acquired from 90 × 90 µm regions. Scale bars, 20 µm. All data are presented as mean ± SEM. Hash symbols indicate statistical significance compared with untreated healthy donor control fibroblasts (# *p* < 0.05, ## *p* < 0.01, ### *p* < 0.001, #### *p* < 0.0001), Asterisks indicate statistical significance compared with untreated patient-derived fibroblasts (* *p* < 0.05, ** *p* < 0.01, *** *p* < 0.001, **** *p* < 0.0001).

Given that mitochondrial fragmentation in patient-derived fibroblasts is associated with increased glucose consumption as a compensatory response to impaired oxidative metabolism [11], we next examined whether restoring mitochondrial architecture was accompanied by metabolic normalization. Consistent with previous reports, LCHADD and VLCADD fibroblasts exhibited significantly elevated glucose consumption under baseline conditions compared with healthy controls (Figure 1C). Treatment with C7 and naloxone reduced glucose uptake on a per-cell-basis, with naloxone showing the strongest reduction and in VLCADD cells even reaching levels observed in control cells (Figure 1C). To determine whether the rescue effect of naloxone was associated with its ROS-scavenging activity, mitochondrial ROS levels were assessed after 72 hours of naloxone treatment. Consistent with the glucose uptake findings, the elevated levels of mitochondrial ROS observed in patient-derived fibroblasts were reduced to levels comparable with the control in the presence of naloxone, indicating that mitochondrial network restoration reverses the increase in ROS (Figure 1D-E).

Since the integrity of the mitochondrial network is governed by a tightly regulated balance between fusion and fission, we analyzed the expression and subcellular distribution of mitochondrial remodeling proteins to evaluate changes associated with naloxone treatment. Under baseline conditions, dynamin-1-like protein 1 (DNM1L) levels were reduced in patient-derived fibroblasts compared with controls, while mitofusin-2 (MFN2) levels showed only minor differences. Following naloxone treatment, DNM1L levels increased in all LCHADD fibroblast lines, accompanied by increased MFN2 expression (Figure 2A-C). Subcellular fractionation revealed that MFN2 was predominantly detected in the mitochondria-enriched fraction, and its abundance in this fraction increased following naloxone treatment (Figure 2D-E). Under baseline conditions, DNM1L was primarily detected in the cytosolic fraction, with reduced levels observed in patient-derived fibroblasts compared with controls. Following naloxone treatment, DNM1L levels increased in the mitochondria-enriched fraction (Figure 2D-E). Consistent with the increased glucose consumption shown in Figure 1C, levels of the major oxidative phosphorylation (OXPHOS) complexes were reduced in all patient-derived fibroblasts (Figure 2D), supporting our previous findings. Following naloxone treatment, OXPHOS protein levels increased in the mitochondria-enriched fraction (Figure 2D-E). These results are in line with the observed reduction in glucose consumption and the restoration of network-like mitochondrial structures (Figure 1A).

**Figure 2.**
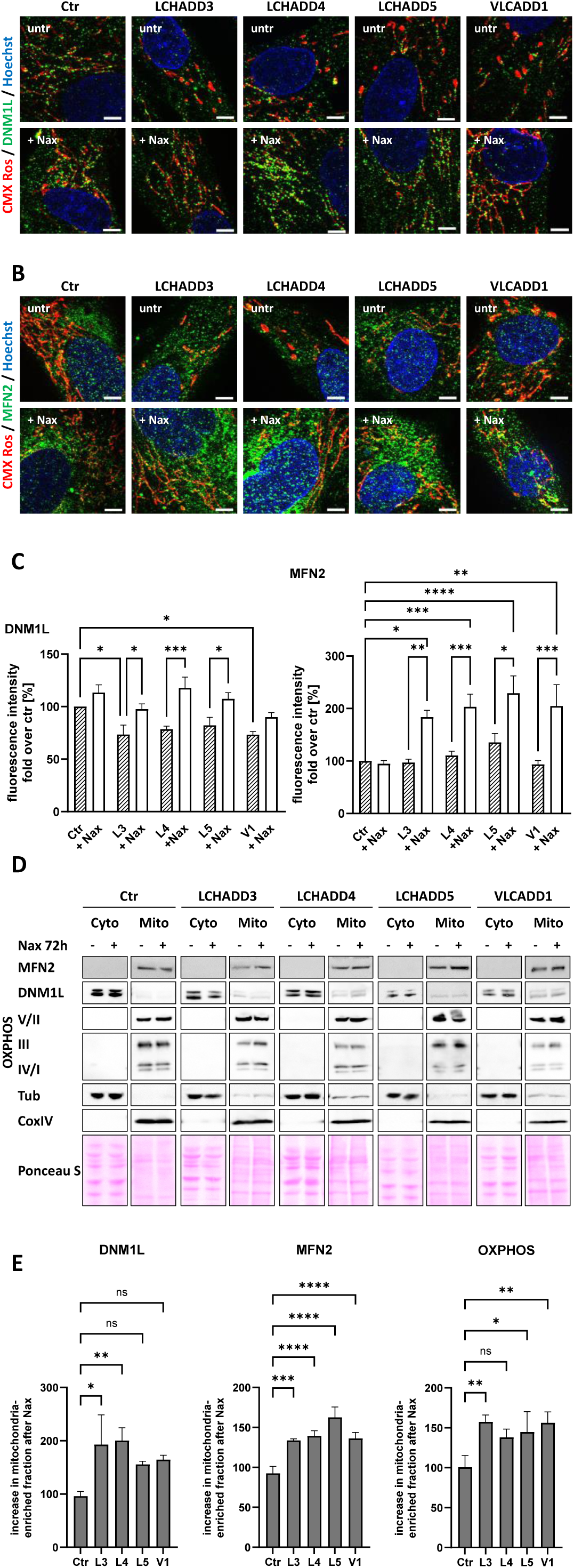
Naloxone alters mitochondrial fusion-fission regulators and OXPHOS in LCHADD and VLCADD patient-derived fibroblasts. **A**, **B** Representative immunofluorescence images of DNM1L (A, green) and MFN2 (B, green) in healthy donor control fibroblasts and patient-derived fibroblasts with or without naloxone (Nax, 1 µM, every 12 h, 72 h). Mitochondria and Nuclei were counterstained with CMXRos (red) and Hoechst (blue), respectively. Scale bars, 5 µm. **C** Quantification of DNM1L (left) and MFN2 (right) fluorescence intensity, expressed as fold change relative to untreated control fibroblasts (≥ 10 cells per condition). **D** Immunoblot analysis of cytosolic (Cyto) and mitochondria-enriched (Mito) fractions from untreated or naloxone-treated fibroblasts (72 h). Fraction purity and equal loading were verified by tubulin (Cyto), COX IV (Mito), and Ponceau S staining. E Quantification of MFN2, DNM1L, and OXPHOS protein abundance, expressed as fold change relative to naloxone-treated healthy donor control fibroblasts. Data from three independent experiments are presented as mean ± SEM. Statistical significance is indicated as: ns, not significant; * *p* < 0.05; ** *p* < 0.01; *** *p* < 0.001; **** *p* < 0.0001.

### Naloxone modulates genes associated with stress fibers and matrix remodeling

Given the pronounced effects of naloxone on mitochondrial network organization, oxidative stress, and metabolic function, RNA sequencing was performed to identify transcriptomic changes associated with naloxone treatment in fibroblasts derived from patients with LCHADD and VLCADD. Differential gene expression analysis revealed distinct transcriptional profiles between healthy controls and untreated patient-derived fibroblasts, as well as between untreated and naloxone-treated patient fibroblasts. Comparison of untreated patient-derived fibroblasts with healthy controls identified 6167 significantly differentially expressed genes (FDR < 0.05), including 3136 upregulated and 3004 downregulated genes. Comparison of untreated and naloxone-treated patient-derived fibroblasts identified 3695 significantly differentially expressed genes (FDR < 0.05), comprising 1764 upregulated and 1931 downregulated genes. In contrast, naloxone treatment induced only minimal transcriptional changes in healthy control fibroblasts, with no significantly differentially expressed genes detected at an FDR threshold of < 0.05 (Table S1). A heatmap of the 50 genes with the highest expression differences demonstrated clear segregation of healthy controls, untreated patient-derived fibroblasts, and naloxone-treated patient-derived fibroblasts (Figure 3A). Volcano plots summarize the distribution of fold changes and statistical significance for both comparisons, highlighting representative differentially expressed genes involved in cytoskeletal organization (ACTC1), fibroblast activation (TGFBI), and extracellular matrix remodeling and production (MMP1, PLAU, COL1A1) (Figure 3B-C).

**Figure 3.**
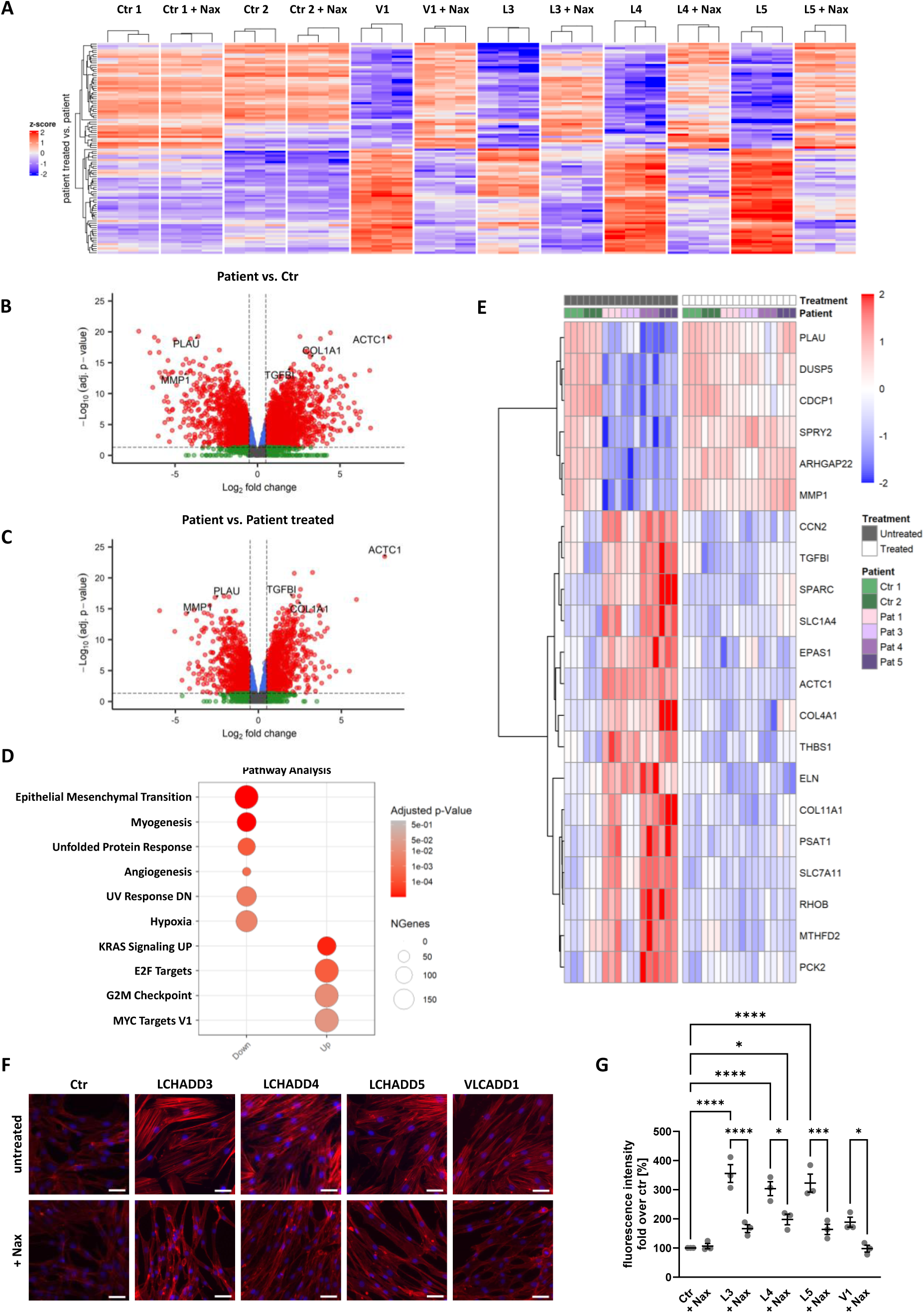
Transcriptomic and cytoskeletal changes in naloxone-treated patient-derived fibroblasts. **A** Heatmap of the top 50 upregulated and top 50 downregulated genes identified by differential gene expression analysis of naloxone-treated versus untreated patient-derived fibroblasts. Gene expression values are displayed as row-wise z-scores. Samples and genes were hierarchically clustered. **B** Volcano plot of differential gene expression between patient-derived fibroblasts and healthy donor control fibroblasts. Gray dots represent genes meeting neither the adjusted P-value (FDR < 0.05) nor the absolute log₂ fold-change (|log₂FC| > 0.5) threshold. Green and blue dots represent genes meeting only the log₂ fold-change or adjusted *P* value threshold, respectively. Red dots represent genes meeting both thresholds. Selected genes (ACTC1, COL1A1, MMP1, PLAU, and TGFBI) are labeled. Positive log₂FC values indicate higher expression in patient-derived fibroblasts. **C** Volcano plot of differential gene expression between naloxone-treated and untreated patient-derived fibroblasts. Colors and thresholds are as described in (B). Positive log₂FC values indicate higher expression in untreated patient-derived fibroblasts. **D** Bubble plot of significantly enriched hallmark pathways in naloxone-treated compared with untreated patient-derived fibroblasts. Pathways enriched among upregulated and downregulated genes are shown separately. Bubble size indicates the number of genes assigned to each pathway, and color represents the adjusted *P* value. **E** Heatmap of selected genes of interest showing differential gene expression in naloxone-treated and untreated patient-derived fibroblasts. Gene expression values are displayed and clustered as in (A). Columns represent individual biological replicates and are annotated by treatment and patient. Red indicates higher expression, whereas blue indicates lower expression. **F** Representative fluorescence images of Phalloidin Atto590-stained control and patient-derived fibroblasts with or without naloxone (Nax; 1 µM, every 12 h). Scale bars, 50 µM. **G** Quantification of F-actin fluorescence intensity, expressed as fold change relative to untreated healthy control fibroblasts (n = 3). Data are presented as mean ± SEM. Statistical significance is indicated as: * *p* < 0.05; ** *p* < 0.01; *** *p* < 0.001; **** *p* < 0.0001.

Differential gene expression analysis revealed significant downregulation of stress-associated pathways following naloxone treatment, including UV response, hypoxia, and the unfolded protein response. Similarly, pathways related to extracellular matrix remodeling and tissue organization, such as epithelial-mesenchymal transition and myogenesis, were also downregulated. In contrast, gene sets associated with cellular activation, survival, and regenerative processes were significantly upregulated (Figure 3D).

To further characterize the transcriptional response to naloxone, individual differentially expressed genes were examined. Naloxone treatment altered the expression of genes associated with cellular stress signaling, metabolic adaptation, cytoskeletal organization, and extracellular matrix remodeling (Figure 3E). Following naloxone treatment, expression of genes involved in stress-associated signaling and negative regulation of MAPK/ERK signaling (DUSP5, SPRY2) was increased. In parallel, expression of genes associated with oxidative and hypoxic stress responses (SLC7A11, EPAS1) and cytoskeletal stress adaptation (RHOB) were reduced. Furthermore, naloxone treatment was associated with transcriptional changes indicative of reduced metabolic stress adaptation in patient-derived fibroblasts. These changes included genes involved in redox homeostasis (SLC7A11), mitochondrial metabolic reprogramming and anaplerotic metabolism (PCK2), mitochondrial one-carbon metabolism (MTHFD2), and serine biosynthesis (PSAT1).

Comparison of healthy control and untreated patient-derived fibroblasts revealed transcriptional changes consistent with altered cytoskeletal organization and extracellular matrix remodeling (Figure 3B). Untreated LCHADD and VLCADD fibroblasts exhibited increased expression of genes associated with actin cytoskeleton organization and stress fiber formation (RHOB, ACTC1). In parallel, genes involved in extracellular matrix production, fibroblast activation, and matrix organization (SPARC, CCN2, COL11A1, COL4A1, ELN, THBS1) were upregulated. Conversely, expression of genes associated with extracellular matrix degradation and cytoskeletal remodeling (MMP1, PLAU, ARHGAP22) was reduced. Taken together, these transcriptional changes define a pro-fibrotic gene expression signature in untreated patient-derived fibroblasts.

Naloxone treatment largely reversed the pro-fibrotic transcriptional profile (Figure 3E). Expression of genes associated with actin cytoskeleton organization and stress fiber formation (RHOB, ACTC1) decreased, while expression of a regulator of actin cytoskeleton dynamics (ARHGAP22) increased. In parallel, expression of genes involved in extracellular matrix production and fibroblast activation (SPARC, TGFBI, CCN2, COL11A1, COL4A1, ELN, THBS1) decreased following naloxone treatment, while expression of genes associated with extracellular matrix degradation and remodeling (MMP1, PLAU) increased.

Since naloxone treatment reversed the expression of genes associated with stress fiber organization and extracellular matrix remodeling, we next investigated whether these transcriptional changes were reflected in altered cytoskeletal architecture. F-actin organization was assessed by phalloidin staining.

Under untreated conditions, patient-derived fibroblasts exhibited prominent phalloidin-positive stress fibers, whereas healthy control fibroblasts displayed less pronounced filamentous actin organization (Figure 3F). Following naloxone treatment, the abundance and organization of stress fibers were markedly altered in all patient-derived fibroblasts. Treated cells displayed thinner and less prominent F-actin bundles, accompanied by a more diffuse intracellular F-actin distribution. Quantitative fluorescence analysis confirmed increased phalloidin signal intensity in untreated patient-derived fibroblasts, which decreased significantly upon naloxone treatment (Figure 3G).

### Development of a 3D bioprinted, vascularized tissue model for LCHADD and VLCADD

Having demonstrated that untreated LCHADD and VLCADD fibroblasts exhibited a pro-fibrotic transcriptional signature accompanied by increased stress fiber formation, and that both features were reversed by naloxone treatment, we next investigated whether these cellular alterations affected vascular network formation in a tissue-like environment. To this end, we established a 3D bioprinted, vascularized tissue model incorporating fibroblasts from healthy donors or patients with LCHADD/VLCADD together with endothelial cells and adipose tissue-derived stem cells (ASCs). In this model, fibroblasts provide structural and biochemical support for vascular network formation through extracellular matrix remodeling and stromal signaling. This enables the assessment of fibroblast-dependent angiogenesis under defined three-dimensional conditions.

The matrix composition was optimized to support vascular network formation. Rheological analysis of hydrogels with varying concentrations identified a formulation with a shear storage modulus of approximately 1400 Pa (4% GelMA) as optimal for network development, closely matching the reported endothelial tissue stiffness [30]. Under these conditions, robust and reproducible vascular structures were formed, whereas increased matrix stiffness impaired network formation (Figure 4A). Bioprinting parameters were subsequently adjusted to ensure structural stability and reproducibility of the constructs. Under the optimized conditions, homogeneous, and mechanically stable tissue constructs that supported vascular network formation were consistently obtained, whereas suboptimal settings resulted in structural instability (Figure 4B). Figure 4C summarizes the overall fabrication workflow of the 3D bioprinted, vascularized tissue constructs. Using these optimized conditions, we observed spontaneous formation of interconnected vascular networks over time. Vascularization was quantified using AngioTool [31], which enabled assessment of network architecture across experimental conditions (Figure 4D).

**Figure 4.**
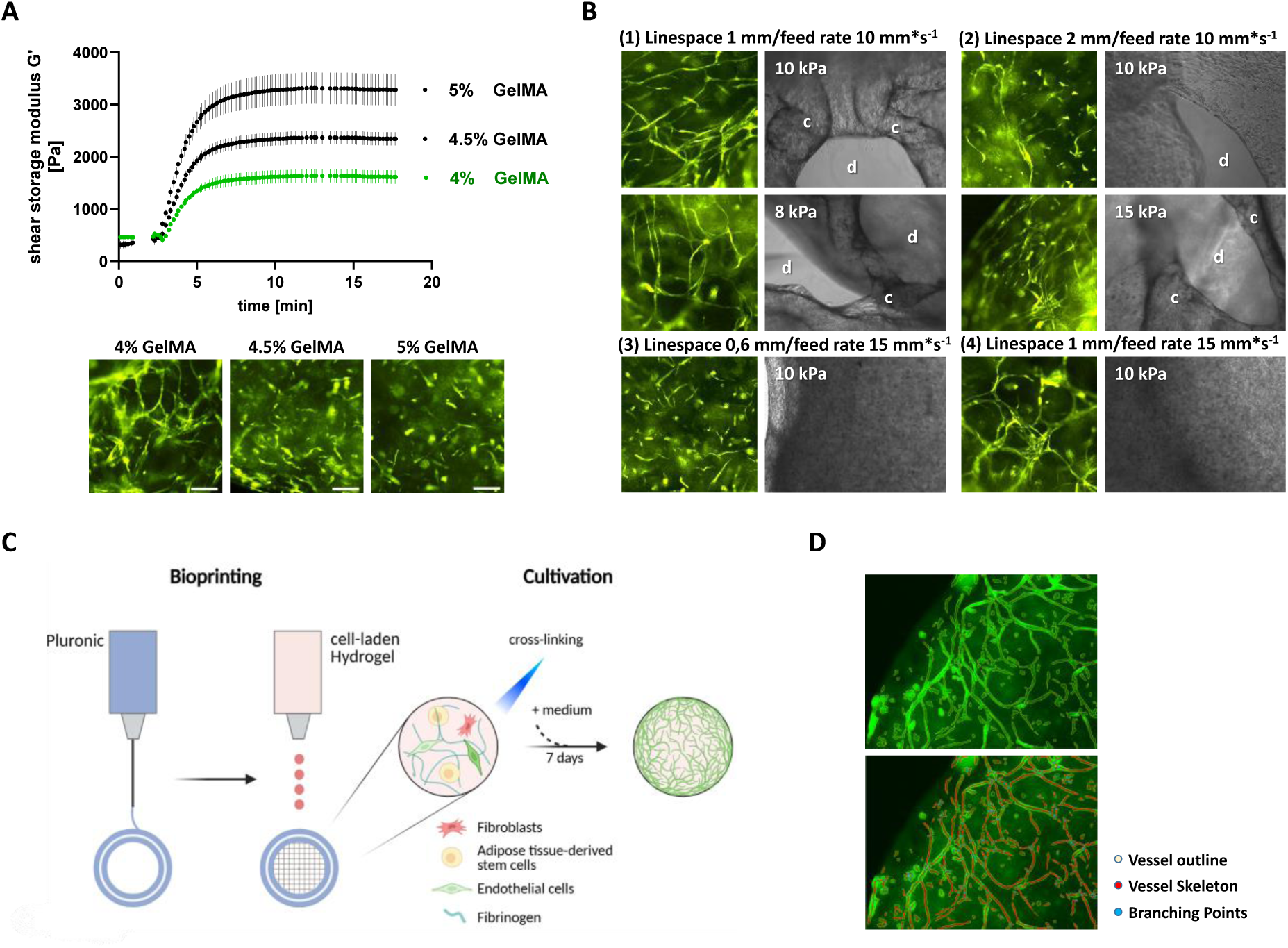
Establishment of a reproducible fully 3D bioprinted vascularized tissue model for LCHADD and VLCADD. **A** Rheological analysis of GelMA bioinks with different concentrations (4%, 4.5%, and 5% w/v). Shear storage modulus (G′) was measured over time. Representative fluorescence images show vascular network formation in 3D bioprinted constructs cultured for 7 days. Scale bars, 200 µm. Data are presented as mean ± SEM (n = 3-5 independent experiments per condition). **B** Evaluation of bioprinting parameters. Printing pressure, feed rate, and infill grid spacing were varied as indicated. Brightfield images show construct morphology, including collapsed (c) and disrupted (d) structures, with corresponding fluorescence images of vascular networks at day 7. **C** Schematic overview of the 3D bioprinting workflow. Thermosensitive Pluronic F-127 was used as a temporary support structure. Cell-laden GelMA was printed and cross-linked by blue-light exposure (390-400 nm). Constructs contained healthy donor control or patient-derived fibroblasts, endothelial cells, and adipose tissue-derived stem cells. **D** Quantification of vascular network formation using AngioTool. The upper panel shows vessel outlines after parameter adjustment; the lower panel shows the analyzed image with vessel outlines (yellow), vessel skeleton (red), and branching points (blue).

After developing a reproducible 3D bioprinted vascularized tissue model, we assessed the functional consequences of the pro-fibrotic fibroblast phenotype on vascular network formation. Vascular network formation was analyzed in 3D constructs containing either fibroblasts from healthy donors or patient-derived fibroblasts. As shown in Figure 5A, constructs containing healthy donor fibroblasts formed interconnected vascular networks within seven days. In contrast, vascularization was consistently impaired in constructs containing LCHADD or VLCADD fibroblasts. Total vessel length was reduced by approximately 30-40% compared with healthy controls and vascular junction density decreased by approximately 30-50%, depending on the patient-derived fibroblast line. These differences were evident by day 5 and became more pronounced by day 7 (Figure 5A-B).

**Figure 5.**
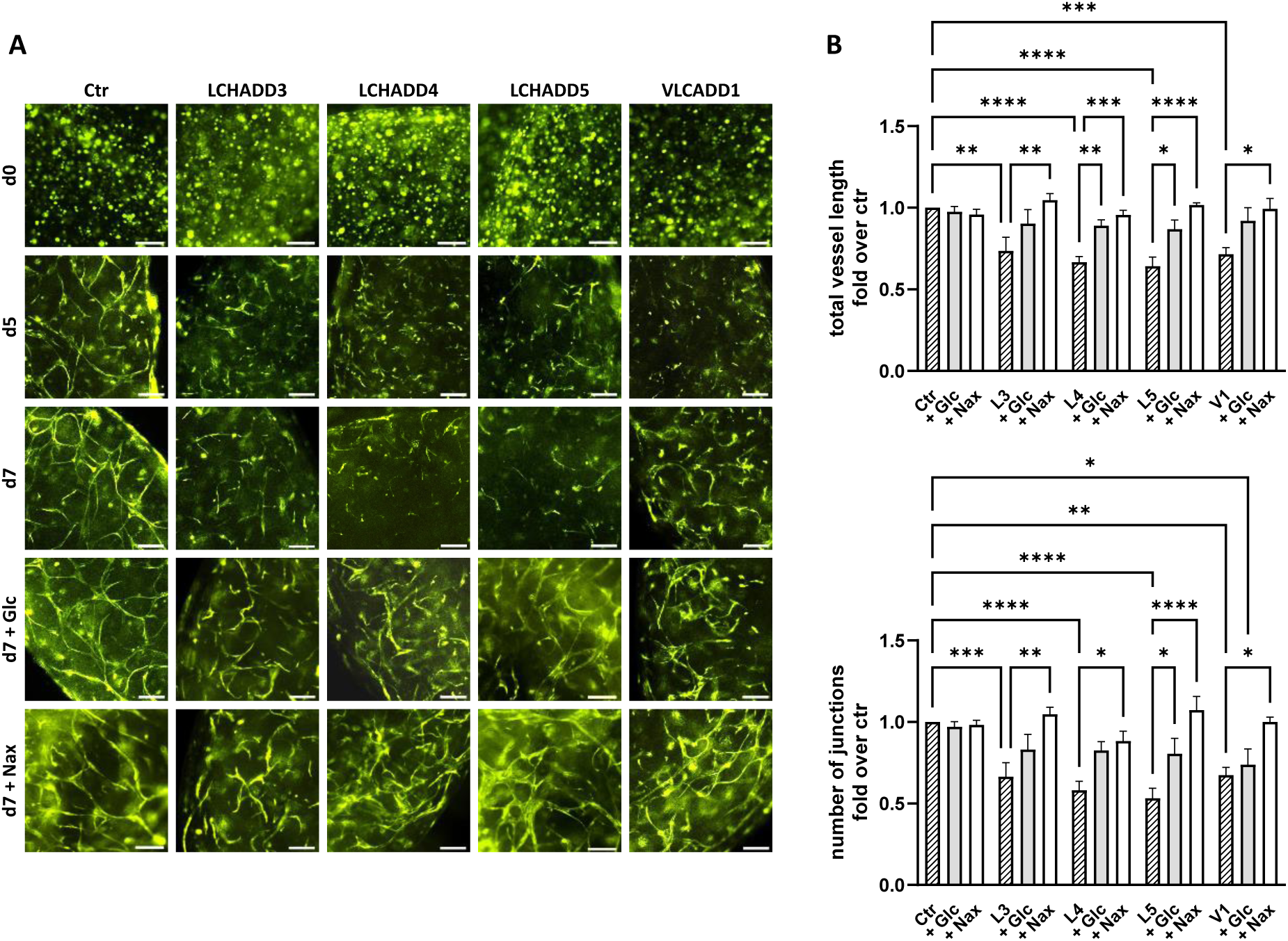
Vascular network formation in 3D bioprinted tissue models containing LCHADD and VLCADD fibroblasts. **A** Representative fluorescence images of 3D bioprinted tissue constructs composed of healthy donor control or patient-derived fibroblasts, human dermal microvascular endothelial cells (HDMVECs; EYFP-labeled), and adipose tissue-derived stem cells (ASCs). Constructs were cultured for 7 days without or with naloxone (Nax; 1 µM, every 12 h) or glucose (Glc; 4.5 g/L). Scale bars, 200 µm. **B** Quantification of vascular network formation, including total vessel length (upper panel) and number of junctions (lower panel), expressed as fold change relative to naloxone-treated healthy donor control fibroblast-containing constructs (n = 3-5 independent experiments per condition, with each experiment comprising two individual tissue surrogates. Data are presented as mean ± SEM. Statistical significance is indicated as: * *p* < 0.05; ** *p* < 0.01; *** *p* < 0.001; **** *p* < 0.0001.

Since naloxone treatment largely reversed the pro-fibrotic transcriptional signature and normalized F-actin organization in patient-derived fibroblasts, its effect on vascular network formation was examined. Glucose supplementation and naloxone treatment did not alter vascular network formation in constructs containing healthy fibroblasts. In contrast, both treatments significantly improved vascularization in constructs containing patient-derived fibroblasts (Figure 5A). Quantitative analysis showed that glucose supplementation increased total vessel length from 60-70% of control values in untreated conditions to 80-90%, whereas naloxone restored vessel length to values comparable to those of healthy controls (Figure 5B). Similarly, vascular junction density increased from 50-70% of control values in untreated constructs to 70-80% with glucose supplementation and to 90-100% following naloxone treatment (Figure 5B). Overall, naloxone showed the most pronounced improvement in vascular network formation in constructs containing LCHADD or VLCADD fibroblasts.

## Discussion

### Redox modulation restores mitochondrial structure in LCHADD and VLCADD fibroblasts

In a previous study, we identified an abnormal mitochondrial phenotype in LCHADD and VLCADD fibroblasts, characterized by fragmented mitochondrial networks, impaired bioenergetic function, and increased oxidative stress [11]. In the present work, we demonstrated that patient-derived fibroblasts consistently exhibited reduced mitochondrial connectivity, metabolic dysfunction, and stress-associated expression profiles. We therefore investigated whether this disease-associated phenotype is pharmacologically reversible. We identified a therapeutic strategy targeting the underlying mechanism of ROS production. Finally, we developed a 3D bioprinted model to study disease-associated effects in a multicellular tissue equivalent.

In Figure 1 we demonstrate that metabolic interventions currently used in clinical management, such as supplementation with glucose, linoleic acid, and triheptanoin (C7), partially improved mitochondrial morphology and function. Carbohydrate support is essential for preventing hypoglycemia and suppressing lipolysis in patients with LCHADD and VLCADD. However, the glucose concentration used in this study represents more an emergency treatment during acute metabolic crises rather than the recommended daily carbohydrate intake in these patients [32, 33]. Linoleic acid is part of the dietary management to avoid the risk of essential fatty acid deficiency and is supplied in small, controlled amounts of specific oils, such as walnut oil [29]. Triheptanoin is a pharmaceutical, medium odd-chain (C7) triglyceride that is used in the therapy for LC-FAOD to stabilize energy production, by providing anaplerotic substrates that can bypass long-chain oxidation [34]. However, all these metabolic interventions appear to be largely compensatory because they primarily bypass the metabolic block in long-chain fatty acid oxidation rather than directly addressing the underlying cellular dysfunction [7, 35, 36]. For this reason, pharmacological targeting of redox imbalance has emerged as a more effective approach. We demonstrated that inhibiting NOX2-dependent ROS production using naloxone resulted in nearly completely restored mitochondrial network integrity and normalized metabolic parameters. These findings support the concept that oxidative stress is a central driver of the disease-associated cellular phenotype in LCHADD and VLCADD, as previously demonstrated by us and others [11, 37].

To further define how the mitochondrial network was restored, we analyzed key regulators of mitochondrial dynamics, focusing on the fission mediator DNM1L and the fusion protein MFN2 (Fig. 2). Previously, we demonstrated that patient-derived fibroblasts exhibit an altered balance of these proteins, consistent with dysregulated mitochondrial dynamics [11]. In the present study, naloxone treatment increased both DNM1L and MFN2 abundance in the mitochondria-enriched fraction, while mitochondrial networks became more elongated and interconnected. At first, this appears contradictory, as increased DNM1L is usually linked to increased mitochondrial fission. However, mitochondrial shape reflects the coordinated activity of the fusion-fission machinery, rather than absolute protein abundance [38]. Therefore, the rounded, clustered mitochondria observed in LCHADD and VLCADD fibroblasts are unlikely to reflect excessive fission alone, but rather a state of impaired mitochondrial dynamics. This may involve reduced fusion competence, defective DNM1L turnover, or insufficient mitophagic clearance. In this scenario, mitochondria fail to remodel dynamically, leading to an accumulation of poorly interconnected, dysfunctional organelles.

Thus, the naloxone-induced upregulation of both DNM1L and MFN2 likely reflects restored mitochondrial plasticity rather than a shift toward either fusion or fission. The increased abundance of DNM1L in the mitochondria-enriched fraction suggests its enhanced recruitment to mitochondria, enabling selective fission events required for segregation and removal of damaged mitochondrial components [39, 40]. In parallel, elevated MFN2 levels may promote outer mitochondrial membrane fusion and stabilize ER-mitochondria contact sites, thereby supporting metabolic coupling and network reintegration [41–43]. Together, these findings support a model in which mitochondrial dysfunction in LCHADD and VLCADD fibroblasts is driven by a loss of dynamic equilibrium rather than a unidirectional imbalance. Restoring both fusion and fission capacity appears essential for reestablishing mitochondrial network integrity and function.

### Naloxone reverses a stress-associated pro-fibrotic fibroblast signature

To investigate the mechanisms underlying the naloxone-induced phenotypic improvement, we next analyzed gene expressions by RNA-seq in patient-derived fibroblasts. Untreated patient fibroblasts showed upregulation of stress-response pathways, as well as gene expression patterns consistent with extracellular matrix production, cytoskeletal remodeling, and fibroblast activation. Naloxone treatment partially normalized these transcriptional signatures (Fig. 3). Notably, in control fibroblasts naloxone had minimal transcriptional impact, suggesting disease state dependent effects and limited off-target transcriptional effects.

In untreated, patient-derived fibroblasts, gene expression indicated a stress-associated pro-fibrotic stromal-like program. This profile included higher expression of extracellular matrix components (ELN, COL4A1) and fibroblast activation/fibrosis markers, and extracellular matrix remodeling (SPARC, CCN2, THBS1, COL11A1) as well as lower expression of matrix-remodeling genes (MMP1, PLAU). These changes have been linked to fibroblast remodeling and fibrotic responses [44–47]. Naloxone partially normalized this signature by lowering expression of genes involved in extracellular matrix production and fibroblast activation, and by restoring the expression of matrix-remodeling genes. These findings suggest that redox modulation affects not only mitochondrial parameters but also promotes a shift toward a less activated fibroblast state.

Untreated LCHADD and VLCADD fibroblasts showed higher RHOB expression, a Rho GTPase regulator involved in stress fiber assembly, and increased ACTC1, which contributes to actin remodeling and contractility. In contrast, ARHGAP22, a negative regulator of Rho/Rac-dependent actin remodeling, was reduced. These transcriptional changes were consistent with phalloidin staining, which revealed prominent stress fibers in patient-derived fibroblasts. Elevated ROS levels have been shown to promote actin polymerization [48] through redox-sensitive Rho GTPase signaling, reinforcing cytoskeletal tension, and myofibroblast activation [49, 50]. At endoplasmic reticulum-mitochondria contact sites, actin polymerization promotes DNM1L recruitment and oligomerization at mitochondrial constriction sites, thereby facilitating mitochondrial fission [51]. Persistent activation of the actin-DNM1L axis during cellular stress is associated with excessive mitochondrial fragmentation, cellular senescence, and tissue dysfunction [52]. Following naloxone treatment, RHOB and ACTC1 expression decreased, whereas ARHGAP22 increased. This was accompanied by fewer stress fibers (Fig. 3) and improved mitochondrial network connectivity. Together, these findings support functional crosstalk between cytoskeletal remodeling and mitochondrial dynamics and indicate that naloxone attenuates both the transcriptional and structural features of fibroblast activation.

### Reversal of fibroblast dysfunction restores vascular network formation in 3D

Our 3D bioprinted, vascularized tissue model provides a human platform to analyze fibroblast-driven vessel formation under defined biomechanical and architectural conditions (Fig. 4). This allows us to test whether patient-derived fibroblasts impair vascular self-organization and whether glucose supplementation or naloxone treatment can restore angiogenic support functions. The present study demonstrates an extension from cell-autonomous 2D phenotypes to structured 3D tissue level function by combining patient-derived fibroblasts with human endothelial cells and adipose tissue-derived stem cells, in a 3D bioprinted vascularized tissue model. Fibroblasts regulate extracellular matrix remodeling, vessel stabilization, and endothelial network formation through mechanical and paracrine interactions with endothelial cells [53, 54]. Constructs with LCHADD or VLCADD fibroblasts showed reduced vessel length and junction density, indicating compromised stromal support at the tissue level (Fig. 5). Both glucose supplementation and naloxone significantly improved vascularization in constructs containing patient-derived fibroblasts, with minimal effects in healthy control constructs, consistent with a disease-specific response. Naloxone produced the strongest effect, consistently restoring vascular network formation toward control levels.

As described above, patient-derived fibroblasts exhibited elevated ROS levels (Fig. 1) and prominent stress fibers (Fig. 3), indicating ROS-dependent cytoskeletal remodeling and an activated fibroblast phenotype. Persistent stress fibers are a hallmark of activated myofibroblasts and have been associated with increased matrix stiffness, altered extracellular matrix turnover, and impaired angiogenesis [55–57]. Thus, naloxone-mediated reductions in ROS and stress fiber abundance may contribute to the improved stromal support for vascular network formation observed in the 3D tissue model. Since naloxone attenuated the pro-fibrotic transcriptional signature in LCHADD and VLCADD fibroblasts, these findings suggest that redox modulation improves cellular function and shifts the microenvironment toward a state more supportive of endothelial self-organization and vascular network formation.

These observations expand the effects of naloxone beyond mitochondrial rescue to include improved fibroblast support at the tissue level, suggesting potential for therapeutic modulation during LCHADD/VLCADD crises. More broadly, our findings imply that chronic oxidative stress in LCHADD and VLCADD leads to maladaptive stromal remodeling, which extends beyond mitochondrial dysfunction. This concept may have broader pathophysiological relevance, as fibroblast activation and extracellular matrix remodeling have been associated with long-term complications of LCHADD, including progressive fibrotic remodeling in cardiomyopathy [37] and retinal changes such as degeneration, pigment alterations, thinning, and subretinal fibrosis or neovascularization [58, 59]. This bioprinted, multicellular 3D model described here provides a screening platform for preclinical assessment of candidate interventions, with the potential to prioritize therapies for subsequent in vivo testing. Although this study was performed in dermal fibroblasts, the identified stress-associated, pro-fibrotic fibroblast phenotype may represent a broader disease mechanism that contributes to tissue remodeling in affected organs.

In conclusion, our findings indicate that oxidative stress is a central driver of a stress-associated, pro-fibrotic fibroblast phenotype in LCHADD and VLCADD that is amenable to pharmacological modulation. The NOX2-inhibitor naloxone attenuated disease-associated abnormalities across molecular, cellular, and tissue levels, including improved mitochondrial integrity, mitigation of the pro-fibrotic transcriptional signature, normalized cytoskeletal organization, and rescued vascular network formation. These findings highlight redox modulation as a mechanistically targeted therapeutic strategy for LCHADD and VLCADD.

## Abbreviations

ASCs: adipose tissue-derived stem cells
C7: Triheptanoin/UX007
Cyto: cytoplasm fraction
DNM1L: dynamine-related protein 1
GelMA: gelatin-methacrylate
Glc: glucose
HDMVECs: human dermal microvascular endothelial cells
HUVECs: human umbilical vein endothelial cells
LA: linoleic acid
LAP: lithium phenyl-2,4,6-trimethylbenzoylphosphinate
LC-FAOD: long-chain fatty acid oxidation disorder
LCHADD: long-chain 3-hydroxyacyl-CoA dehydrogenase deficiency
MFN2: mitofusin 2
Mito: mitochondria-enriched fraction
Nax: naloxone
NOX2: NADPH oxidase 2
OXPHOS: oxidative phosphorylation
ROS: reactive oxygen species
TNBS: 2,4,6-trinitrobenzene sulfonic acid
VLCADD: very-long-chain acyl-CoA dehydrogenase deficiency

## Acknowledgement

We thank Mag. Verena Jeller and Nora Kaiser, PhD for technical support. Funding was received from the Austrian Science Fund https://doi.org/10.55776/FG15 and 10.55776/PAT1161725, the Federal Ministry Republic of Austria for Education, Science and Research (Project “Replacement of animal experiments in science”) the Tiroler Wissenschaftsförderung (“Gefördert aus Mitteln des Landes Tirol”) and the “Tirol-Kliniken GmbH”.

## Competing interests

The authors have declared that no competing interests exist.

## Declarations

During the preparation of this manuscript, the authors used the university-provided Academic AI and DeepL exclusively for grammar and language editing. The authors reviewed and approved all revisions and take full responsibility for the final content of the manuscript.

Schematic figures were created using BioRender.com. The authors confirm that they have the appropriate license to use BioRender for publication and that the content is original and created by the authors.

## Notes

### Competing Interest Statement

The authors have declared no competing interest.

### Summary of Updates

We have updated Figure 5A since there was an image duplication by mistake.

