## Supplemental Data for "Naloxone as mitochondrial phenotype rescuer in a 3D bioprinted LCHADD/VLCADD model"

This file includes:

Figures S1 to S4

Tables S1 to S5

### Supplementary Figures

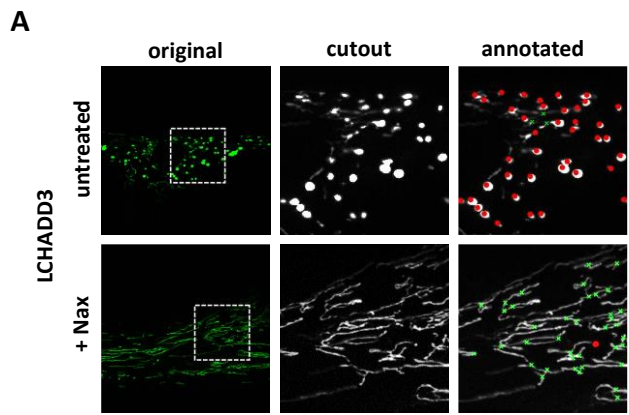

**Figure S1:** Representative MiMoQuant analysis of mitochondrial morphology. **A** Representative images of LCHADD patient-derived fibroblasts treated with naloxone (1 μM, every 12 h) for up to 72 h. Mitochondrial branches and dots were quantified automatically using MiMoQuant and are shown in green and red, respectively.

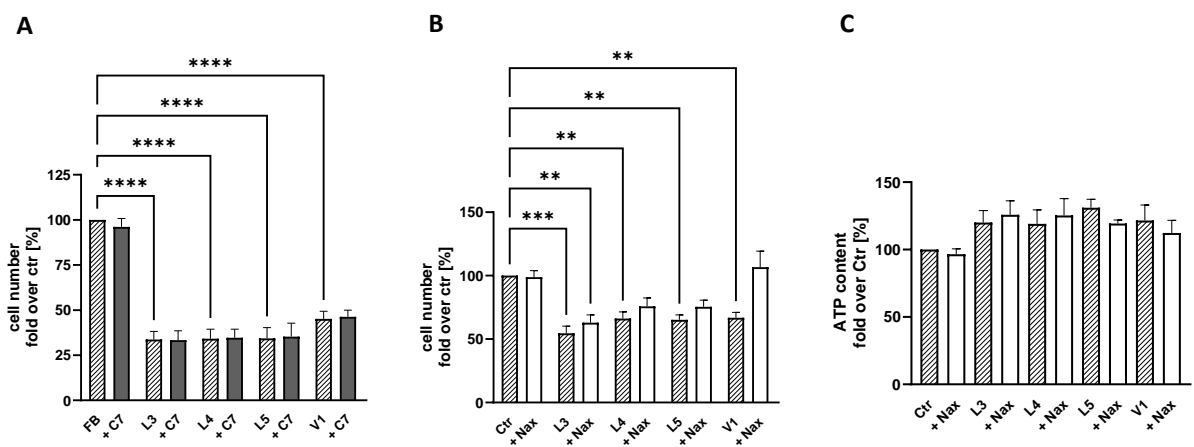

**Figure S2.** Effects of C7 and naloxone on cell number and ATP levels in LCHADD and VLCADD patient-derived fibroblasts. **A, B** Cell number of control and patient-derived fibroblasts under untreated conditions or after 72 h treatment with C7 (**A**; 2.5 mmol/L; $n = 5$) or naloxone (**B**; Nax; 1 μM, every 12 h; $n = 4$). **C** Intracellular ATP content (CellTiter-Glo<sup>®</sup> Luminescent Cell Viability Assay (Promega, Mannheim, Germany)) in untreated or naloxone-treated fibroblasts after 72 h ($n = 5$). Data are expressed as fold change relative to untreated control healthy donor cells and presented as mean ± SEM. Statistical significance is indicated as: \*\* $p < 0.01$; \*\*\* $p < 0.001$; \*\*\*\* $p < 0.0001$.

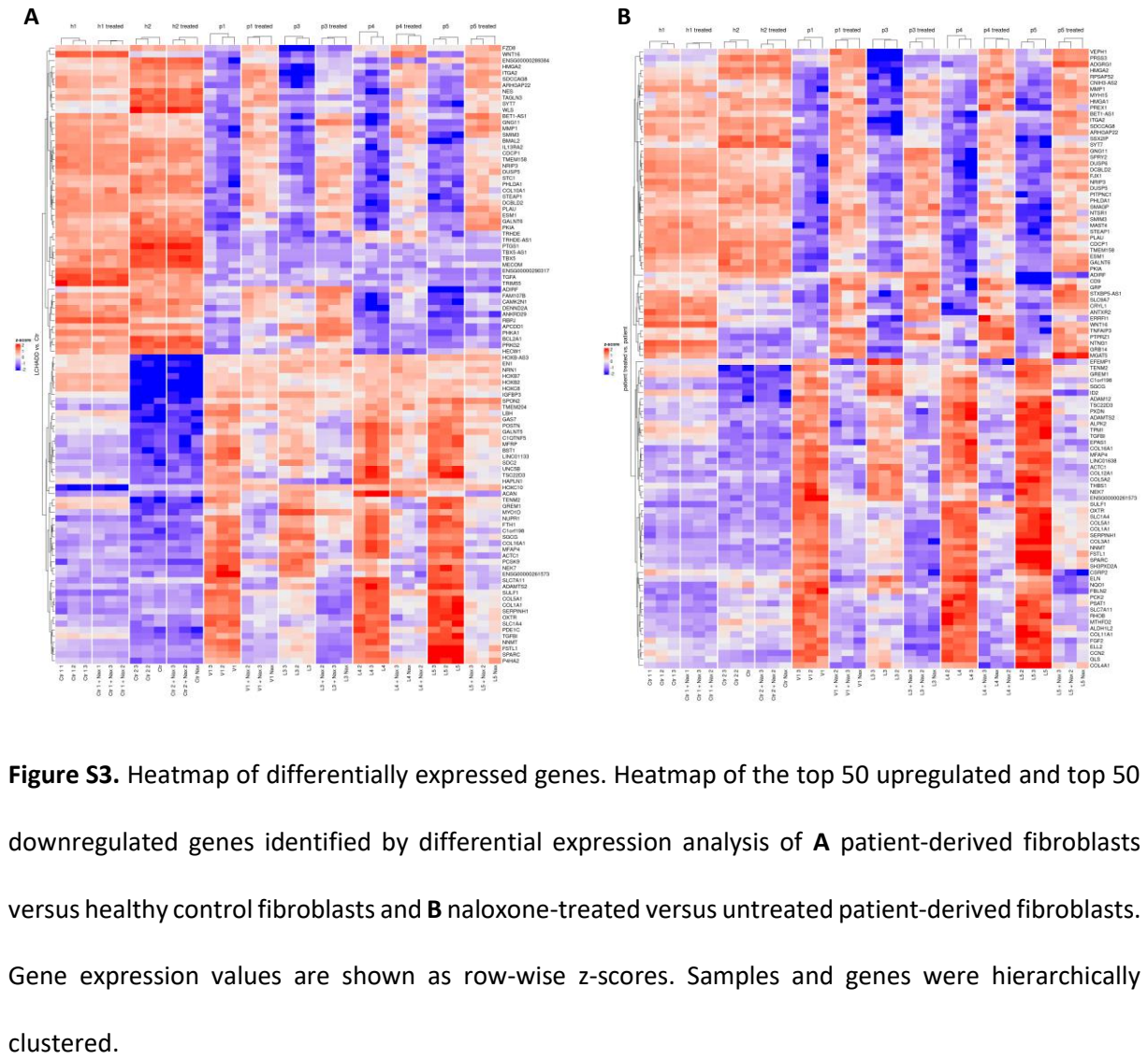

Figure S3. Heatmap of differentially expressed genes. Heatmap of the top 50 upregulated and top 50 downregulated genes identified by differential expression analysis of A patient-derived fibroblasts versus healthy control fibroblasts and B naloxone-treated versus untreated patient-derived fibroblasts. Gene expression values are shown as row-wise z-scores. Samples and genes were hierarchically clustered.

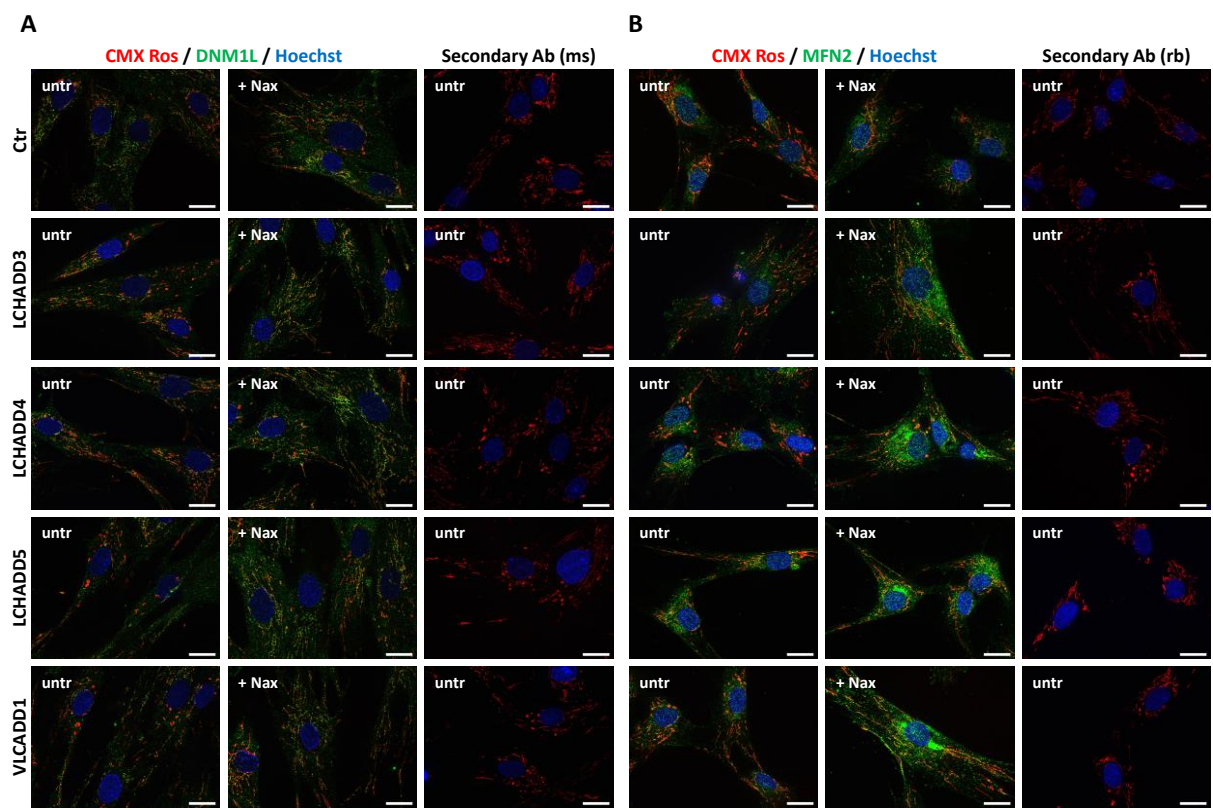

**Figure S4.** Uncropped immunofluorescence images related to Fig. 2. (A, B) Representative merged immunofluorescence images of DNM1L (A, green) and MFN2 (B, green) in control and patient-derived fibroblasts with or without naloxone (Nax, 1 μM, every 12 h, 72 h). Mitochondria and Nuclei were counterstained with CMXRos (red) and Hoechst (blue), respectively. Secondary antibody controls are shown for DNM1L (mouse) and MFN2 (rabbit). Scale bars, 5 μm.

**Table S1.** Summary of differential gene expression and pathway analysis. Overview of significantly up- and downregulated genes and enriched pathways identified by differential expression (DE) analysis.

| DE_genes | Down | NotSig | Up |
| --- | --- | --- | --- |
| Ctr + Nax vs. Ctr | 0 | 14011 | 0 |
| patient treated vs. patient | 1931 | 10316 | 1764 |
| LCHADD vs. Ctr | 3004 | 7844 | 3163 |
| LCHADD + Nax vs. Ctr | 1563 | 10664 | 1784 |
| LCHADD vs. Ctr + Nax | 2631 | 8586 | 2794 |
| LCHADD + Nax vs. Ctr + Nax | 834 | 12062 | 1115 |
| Ctr1 + Nax vs. Ctr1 | 0 | 14011 | 0 |
| Ctr2 + Nax vs. Ctr2 | 0 | 14011 | 0 |
| V1 + Nax vs. V1 | 649 | 12877 | 485 |
| L3 + Nax vs. L3 | 440 | 13167 | 404 |
| L4 + Nax vs. L4 | 614 | 12946 | 451 |
| L5 + Nax vs. L5 | 886 | 12442 | 683 |
| **DE_pathways** | **Down** | **NotSig** | **Up** |
| Ctrl + Nax vs. Ctrl | 0 | 14011 | 0 |
| patient treated vs. patient | 1931 | 10316 | 1764 |
| LCHADD vs. Ctrl | 3004 | 7844 | 3163 |
| LCHADD + Nax vs. Ctrl | 1563 | 10664 | 1784 |
| LCHADD vs. Ctrl + Nax | 2631 | 8586 | 2794 |
| LCHADD + Nax vs. Ctrl + Nax | 834 | 12062 | 1115 |
| Ctrl1 + Nax vs. Ctrl1 | 0 | 14011 | 0 |
| Ctrl2 + Nax vs. Ctrl2 | 0 | 14011 | 0 |
| V1 + Nax vs. V1 | 649 | 12877 | 485 |
| L3 + Nax vs. L3 | 440 | 13167 | 404 |
| L4 + Nax vs. L4 | 614 | 12946 | 451 |
| L5 + Nax vs. L5 | 886 | 12442 | 683 |

**Table S2.** Differential expression analysis of the top 50 upregulated genes in naloxone-treated versus untreated patient-derived fibroblasts. Differential expression statistics include logFC, logCPM, F-statistic, and FDR-adjusted p-values. Positive logFC values indicate higher expression in the first group of the comparison.

| gene_name | logFC | logCPM | F | FDR | PValue |
| --- | --- | --- | --- | --- | --- |
| ARHGAP22 | 2,06206494 | 6,87142831 | 522,454813 | 9,9417E-18 | 4,967E-21 |
| GNG11 | 1,77444467 | 8,61219227 | 512,078527 | 1,1677E-17 | 6,6676E-21 |
| PLAU | 2,60208877 | 7,15128741 | 499,611588 | 1,4954E-17 | 9,6055E-21 |
| PKIA | 2,92081049 | 4,61024981 | 402,810717 | 2,848E-16 | 2,4392E-19 |
| TMEM158 | 3,87955225 | 8,49160069 | 345,393306 | 1,5066E-15 | 2,043E-18 |
| NTSR1 | 5,94417059 | 4,24325635 | 321,411037 | 2,0911E-15 | 3,2834E-18 |
| SMAGP | 1,37763319 | 6,38897361 | 334,645844 | 2,0911E-15 | 3,2047E-18 |
| SMIM3 | 1,94020645 | 6,58526409 | 325,565498 | 2,5582E-15 | 4,7472E-18 |
| DCBLD2 | 1,52839303 | 8,00459415 | 321,898202 | 2,7895E-15 | 5,5745E-18 |
| SYT7 | 3,26331316 | 3,07269291 | 268,649377 | 4,7348E-15 | 1,0476E-17 |
| MMP1 | 4,35240811 | 10,3733915 | 300,595737 | 6,4551E-15 | 1,4743E-17 |
| ADGRG1 | 3,03507769 | 3,72678069 | 291,678937 | 6,6832E-15 | 1,6112E-17 |
| ITGA2 | 2,81612541 | 7,24546569 | 280,920108 | 1,4163E-14 | 3,8412E-17 |
| NTNG1 | 1,80838011 | 5,63656469 | 264,042436 | 3,1322E-14 | 9,1658E-17 |
| CRYL1 | 1,74494691 | 4,5113713 | 260,701302 | 3,661E-14 | 1,0992E-16 |
| SPRY2 | 2,25039261 | 5,78440912 | 255,990712 | 4,3892E-14 | 1,4115E-16 |
| HMGA1 | 2,10974928 | 9,94012972 | 256,004155 | 4,3892E-14 | 1,4074E-16 |
| HMGA2 | 1,6776735 | 6,90182036 | 255,188075 | 4,3892E-14 | 1,4724E-16 |
| CDCP1 | 3,71441924 | 4,96049594 | 238,961157 | 8,4395E-14 | 3,1924E-16 |
| FJX1 | 1,70274686 | 5,53139244 | 228,42777 | 1,6156E-13 | 6,8384E-16 |
| ANTXR2 | 0,96649843 | 7,09129097 | 223,98676 | 2,0537E-13 | 8,9413E-16 |
| PREX1 | 2,22014427 | 3,77861572 | 223,581945 | 2,0981E-13 | 9,5839E-16 |
| GALNT6 | 1,4557129 | 7,17092606 | 222,506646 | 2,1114E-13 | 9,7954E-16 |
| BET1-AS1 | 2,79583026 | 4,11592536 | 222,688241 | 2,2253E-13 | 1,0483E-15 |
| DUSP5 | 1,71666525 | 4,97442683 | 220,309993 | 2,3568E-13 | 1,127E-15 |
| NRIP3 | 1,79433671 | 4,83804039 | 219,542765 | 2,4374E-13 | 1,1829E-15 |
| SDCCAG8 | 1,02437757 | 6,1151062 | 211,655839 | 3,8316E-13 | 1,9417E-15 |
| VEPH1 | 1,2415909 | 5,80608194 | 208,960035 | 4,3218E-13 | 2,3134E-15 |
| STEAP1 | 2,00350357 | 6,48178603 | 201,925186 | 6,7038E-13 | 3,6842E-15 |
| ESM1 | 3,2480906 | 5,13934285 | 192,1771 | 1,2771E-12 | 7,5656E-15 |
| SSX2IP | 1,55250368 | 5,78373116 | 189,226639 | 1,461E-12 | 8,8636E-15 |
| WNT16 | 3,14976844 | 5,27741099 | 188,671577 | 1,6113E-12 | 9,8904E-15 |
| ADIRF | 1,90096391 | 10,4163093 | 187,188932 | 1,6462E-12 | 1,0233E-14 |
| MAST4 | 1,59618448 | 4,88236137 | 187,090222 | 1,6462E-12 | 1,0339E-14 |
| SLC9A7 | 1,35422993 | 5,09198553 | 184,323909 | 1,9624E-12 | 1,2606E-14 |
| DUSP6 | 2,68565775 | 6,27200289 | 181,506836 | 2,3585E-12 | 1,5487E-14 |
| PITPNC1 | 1,74645926 | 4,37598765 | 181,185684 | 2,4033E-12 | 1,5953E-14 |
| STXBP5-AS1 | 2,09458363 | 3,42748247 | 180,892291 | 2,4885E-12 | 1,6695E-14 |
| PHLDA1 | 2,11248818 | 7,52033282 | 179,738786 | 2,5838E-12 | 1,7623E-14 |
| CNIH3-AS2 | 1,7228773 | 3,16905226 | 179,711811 | 2,5935E-12 | 1,814E-14 |
| CD9 | 1,43879424 | 7,46400064 | 179,465636 | 2,5935E-12 | 1,7981E-14 |
| PTPRZ1 | 3,40776024 | 1,37689023 | 148,291441 | 3,0511E-12 | 2,1994E-14 |
| TNFAIP3 | 1,40243981 | 4,53157246 | 176,912337 | 3,0511E-12 | 2,1841E-14 |
| PRSS3 | 2,46685449 | 5,84598157 | 174,885803 | 3,2792E-12 | 2,4575E-14 |
| MYH15 | 5,03497162 | 1,27064412 | 143,45519 | 4,818E-12 | 3,7826E-14 |
| RPSAP52 | 1,67118614 | 4,60959377 | 168,432609 | 5,2524E-12 | 4,1986E-14 |
| GRB14 | 2,57911493 | 3,18848598 | 165,698806 | 5,6534E-12 | 4,6402E-14 |
| ERRFI1 | 1,83144509 | 6,14630977 | 166,595138 | 5,7947E-12 | 4,8314E-14 |
| MGAT5 | 1,11079487 | 5,56716324 | 166,471533 | 5,7947E-12 | 4,8803E-14 |
| GRP | 3,29435179 | 3,86748024 | 166,631962 | 5,955E-12 | 5,1003E-14 |

**Table S3.** Differential expression analysis of the top 50 downregulated genes in naloxone-treated versus untreated patient-derived fibroblasts. Differential expression statistics include logFC, logCPM, F-statistic, and FDR-adjusted p-values. Negative logFC values indicate lower expression in the first group of the comparison.

| gene_name | logFC | logCPM | F | FDR | PValue |
| --- | --- | --- | --- | --- | --- |
| ACTC1 | -7,616497246 | 4,76049028 | 808,705878 | 3,3838E-24 | 2,4151E-28 |
| SLC7A11 | -3,267043936 | 4,67460352 | 1030,1624 | 1,4132E-21 | 2,0173E-25 |
| SPARC | -2,173893608 | 11,3222415 | 980,500094 | 1,9087E-21 | 4,0868E-25 |
| NNMT | -2,54154457 | 6,59240184 | 647,561808 | 7,2556E-19 | 2,0714E-22 |
| COL5A1 | -2,475998833 | 7,21860568 | 532,153925 | 8,8652E-18 | 3,789E-21 |
| TGFBI | -2,100278509 | 10,1620263 | 532,040108 | 8,8652E-18 | 3,7964E-21 |
| ELN | -5,913694022 | 5,38919659 | 471,694429 | 3,2983E-17 | 2,3541E-20 |
| COL1A1 | -2,382361888 | 12,3808852 | 440,906779 | 7,6035E-17 | 5,9695E-20 |
| SULF1 | -1,969378291 | 8,35444154 | 395,955642 | 3,0712E-16 | 2,8495E-19 |
| EFEMP1 | -3,647660571 | 5,0981444 | 392,230958 | 3,3663E-16 | 3,3637E-19 |
| COL4A1 | -2,779105829 | 5,46572186 | 370,893484 | 6,8996E-16 | 7,3866E-19 |
| FGF2 | -2,056198205 | 6,40041579 | 362,829487 | 8,8055E-16 | 1,0055E-18 |
| ADAMTS2 | -1,461503511 | 6,18929785 | 359,392954 | 9,5031E-16 | 1,153E-18 |
| COL11A1 | -3,872283422 | 3,3712255 | 328,253302 | 1,5066E-15 | 2,0313E-18 |
| RHOB | -1,736462536 | 5,40052803 | 341,835594 | 1,6598E-15 | 2,3693E-18 |
| CCN2 | -3,189765053 | 8,29007928 | 331,037884 | 2,2789E-15 | 3,7409E-18 |
| COL5A2 | -1,909270702 | 7,08452221 | 329,893955 | 2,294E-15 | 3,9296E-18 |
| SERPINH1 | -1,844861458 | 8,53014815 | 326,503513 | 2,5515E-15 | 4,5527E-18 |
| MFAP4 | -2,754584546 | 4,76738244 | 325,059107 | 2,5698E-15 | 4,9522E-18 |
| NEK7 | -1,771952686 | 7,22616962 | 313,419362 | 3,9383E-15 | 8,1515E-18 |
| FSTL1 | -1,224618148 | 9,53911966 | 309,265872 | 4,6003E-15 | 9,8501E-18 |
| THBS1 | -2,233952004 | 7,92450666 | 298,28899 | 6,6832E-15 | 1,6441E-17 |
| TPM1 | -1,555871769 | 9,71958443 | 297,96384 | 6,6832E-15 | 1,6695E-17 |
| ENSG00000261573 | -3,689937794 | 3,60205262 | 246,864954 | 1,2891E-14 | 3,3333E-17 |
| MTHFD2 | -1,84708272 | 5,86874457 | 283,332454 | 1,2891E-14 | 3,4041E-17 |
| PSAT1 | -2,664917431 | 5,81585044 | 273,867773 | 1,9756E-14 | 5,4991E-17 |
| COL16A1 | -1,477230618 | 6,02206187 | 264,598367 | 3,1123E-14 | 8,8852E-17 |
| GREM1 | -1,786351617 | 9,63940061 | 260,17109 | 3,661E-14 | 1,1236E-16 |
| NQO1 | -1,076334889 | 8,39187812 | 255,286855 | 4,3892E-14 | 1,4635E-16 |
| TENM2 | -2,386622432 | 5,62524213 | 250,600366 | 4,5238E-14 | 1,5545E-16 |
| ALPK2 | -2,085678697 | 3,85289305 | 254,148272 | 4,5238E-14 | 1,5821E-16 |
| PXDN | -1,489082984 | 8,18902689 | 249,372385 | 5,6831E-14 | 2,0281E-16 |
| TSC22D3 | -2,231173261 | 4,09746553 | 248,312157 | 5,9895E-14 | 2,2138E-16 |
| EPAS1 | -1,988405288 | 6,40835422 | 247,746436 | 5,9895E-14 | 2,2229E-16 |
| OXTR | -2,120711727 | 6,83653974 | 237,577323 | 1,0313E-13 | 3,9746E-16 |
| C1orf198 | -1,761466803 | 5,62878691 | 234,843916 | 1,1894E-13 | 4,669E-16 |
| COL12A1 | -2,165755367 | 6,99284462 | 234,145024 | 1,2152E-13 | 4,8571E-16 |
| FBLN2 | -1,597238017 | 6,17511613 | 232,729888 | 1,2985E-13 | 5,2827E-16 |
| COL3A1 | -2,285313473 | 8,9754222 | 231,14563 | 1,401E-13 | 5,7994E-16 |
| PCK2 | -1,900693185 | 4,92765499 | 228,30255 | 1,6156E-13 | 6,9188E-16 |
| LINC01638 | -3,731032256 | 1,31225164 | 155,029081 | 2,0726E-13 | 9,3194E-16 |
| ADAM12 | -2,098845506 | 5,88144688 | 223,595855 | 2,0726E-13 | 9,1779E-16 |
| ID2 | -1,298533014 | 7,8939711 | 215,570509 | 3,0682E-13 | 1,511E-15 |
| SLC1A4 | -1,74281387 | 3,79798164 | 212,921427 | 3,6589E-13 | 1,828E-15 |
| SGCG | -2,614721816 | 2,79017951 | 204,605282 | 3,97E-13 | 2,0401E-15 |
| ALDH1L2 | -1,701770318 | 4,89391709 | 209,867518 | 4,139E-13 | 2,186E-15 |
| ELL2 | -1,445513546 | 6,52828693 | 209,885779 | 4,139E-13 | 2,177E-15 |
| CSRP2 | -2,150601318 | 4,24192289 | 205,529547 | 5,3853E-13 | 2,9211E-15 |
| GLS | -1,068441483 | 6,311061 | 201,454998 | 6,8269E-13 | 3,8006E-15 |
| SH3PXD2A | -1,056600157 | 5,42095198 | 198,066433 | 8,4881E-13 | 4,786E-15 |

**Table S4.** Differential expression analysis of the top 50 upregulated genes in patient-derived fibroblasts versus healthy donor control fibroblasts. Differential expression statistics include logFC, logCPM, F-statistic, and FDR-adjusted p-values. Positive logFC values indicate higher expression in the first group of the comparison.

| gene_name | logFC | logCPM | F | FDR | PValue |
| --- | --- | --- | --- | --- | --- |
| HOXC10 | 4,42618435 | 5,95482952 | 884,7312 | 2,0618E-24 | 1,4444E-20 |
| SPARC | 2,57925629 | 11,3222415 | 764,816917 | 1,7195E-23 | 6,0231E-20 |
| ACTC1 | 8,02333261 | 4,76049028 | 440,267856 | 2,424E-23 | 6,7926E-20 |
| SULF1 | 3,82516904 | 8,35444154 | 693,793708 | 7,3954E-23 | 1,4336E-19 |
| NNMT | 3,02798561 | 6,59240184 | 495,127241 | 1,0963E-20 | 1,2801E-17 |
| SLC7A11 | 2,87485596 | 4,67460352 | 489,32024 | 1,3404E-20 | 1,4447E-17 |
| COL5A1 | 3,14395028 | 7,21860568 | 454,421677 | 3,8463E-20 | 3,3681E-17 |
| COL1A1 | 3,19833112 | 12,3808852 | 415,029102 | 1,4395E-19 | 1,1864E-16 |
| HOXB-AS3 | 3,99988545 | 6,29545769 | 364,681566 | 2,7714E-19 | 2,0437E-16 |
| HOXC8 | 2,20583134 | 6,36763579 | 334,698053 | 3,2559E-18 | 1,8247E-15 |
| IGFBP3 | 3,40068262 | 10,7366834 | 329,318928 | 4,0285E-18 | 2,0905E-15 |
| LINC01133 | 3,96882214 | 7,34587792 | 311,04441 | 9,0908E-18 | 3,8597E-15 |
| GAS7 | 3,58341857 | 3,4669512 | 294,078388 | 1,0864E-17 | 4,3492E-15 |
| TGFB1 | 2,04054862 | 10,1620263 | 294,996499 | 1,9233E-17 | 7,2831E-15 |
| HAPLN1 | 3,85744674 | 6,35790213 | 277,373638 | 3,2899E-17 | 1,213E-14 |
| TENM2 | 4,40670463 | 5,62524213 | 279,036578 | 3,4137E-17 | 1,2264E-14 |
| COL16A1 | 2,02998226 | 6,02206187 | 280,818916 | 3,8566E-17 | 1,3179E-14 |
| MFAP4 | 3,66505543 | 4,76738244 | 278,83149 | 4,3394E-17 | 1,4476E-14 |
| SGCG | 6,81119651 | 2,79017951 | 263,483518 | 5,8261E-17 | 1,8984E-14 |
| SERPINH1 | 2,21202097 | 8,53014815 | 265,893763 | 8,2894E-17 | 2,4711E-14 |
| C1orf198 | 2,53088673 | 5,62878691 | 257,490576 | 1,3013E-16 | 3,575E-14 |
| SPON2 | 2,9088032 | 8,52136446 | 253,951903 | 1,5749E-16 | 4,1633E-14 |
| HOXB7 | 3,2128151 | 5,75400012 | 222,246592 | 2,7335E-16 | 6,3833E-14 |
| UNC5B | 3,1880134 | 5,02226387 | 241,556023 | 3,187E-16 | 7,2022E-14 |
| HOXB2 | 2,66537039 | 4,86325219 | 226,209443 | 3,6809E-16 | 8,0584E-14 |
| BST1 | 2,6426089 | 5,093745 | 237,130575 | 4,096E-16 | 8,8291E-14 |
| MFRP | 3,22799416 | 4,50933933 | 236,079601 | 4,4562E-16 | 9,46E-14 |
| ACAN | 5,89923609 | 5,97060767 | 235,840273 | 4,6028E-16 | 9,6254E-14 |
| C1QTNF5 | 3,21026642 | 4,51124679 | 232,654916 | 5,4464E-16 | 1,1059E-13 |
| P4HA2 | 1,49128709 | 7,41332254 | 227,350298 | 7,2844E-16 | 1,458E-13 |
| ENSG00000261573 | 5,02556888 | 3,60205262 | 198,216483 | 8,6961E-16 | 1,6691E-13 |
| FSTL1 | 1,33224396 | 9,53911966 | 221,467242 | 1,0442E-15 | 1,977E-13 |
| FTH1 | 1,92274226 | 11,8268923 | 217,633039 | 1,3264E-15 | 2,478E-13 |
| NEK7 | 1,91737689 | 7,22616962 | 214,462524 | 1,6215E-15 | 2,9126E-13 |
| TSC22D3 | 2,87839792 | 4,09746553 | 214,159651 | 1,6963E-15 | 3,0084E-13 |
| GREM1 | 2,15092529 | 9,63940061 | 213,403548 | 1,7346E-15 | 3,0379E-13 |
| NRN1 | 2,69149675 | 7,01907187 | 211,399572 | 1,993E-15 | 3,4474E-13 |
| NUPR1 | 3,37368125 | 9,09114099 | 209,600283 | 2,2167E-15 | 3,742E-13 |
| PDE1C | 2,15854713 | 4,16957333 | 208,650633 | 2,3917E-15 | 3,9423E-13 |
| PCSK9 | 5,2292221 | 3,32275077 | 215,161806 | 2,5184E-15 | 4,103E-13 |
| SLC1A4 | 2,29893888 | 3,79798164 | 206,52374 | 2,7689E-15 | 4,359E-13 |
| ADAMTS2 | 1,41379514 | 6,18929785 | 205,700109 | 2,8641E-15 | 4,4588E-13 |
| MYO1D | 2,85879227 | 5,16603127 | 200,694215 | 4,0245E-15 | 5,9355E-13 |
| POSTN | 4,44601976 | 9,48120674 | 199,971066 | 4,2031E-15 | 6,1343E-13 |
| OXTR | 2,60600582 | 6,83653974 | 197,686944 | 4,9098E-15 | 7,0918E-13 |
| EN1 | 3,1883188 | 5,14300953 | 193,619312 | 5,819E-15 | 8,0723E-13 |
| LBH | 2,08260978 | 6,20326352 | 194,767788 | 6,0048E-15 | 8,2483E-13 |
| TMEM204 | 2,95608052 | 3,98042236 | 192,242956 | 7,4861E-15 | 1,0085E-12 |
| GALNT5 | 2,71948642 | 6,60504988 | 189,580731 | 8,6398E-15 | 1,1529E-12 |
| SDC2 | 2,12647363 | 6,40035882 | 188,337135 | 9,4294E-15 | 1,2464E-12 |

**Table S5.** Differential expression analysis of the top 50 downregulated genes in patient-derived fibroblasts versus healthy donor control fibroblasts. Differential expression statistics include logFC, logCPM, F-statistic, and FDR-adjusted p-values. Negative logFC values indicate lower expression in the first group of the comparison.

| gene_name | logFC | logCPM | F | FDR | PValue |
| --- | --- | --- | --- | --- | --- |
| TRIM55 | -7,19751044 | 3,63767159 | 505,074961 | 5,7788E-25 | 8,0967E-21 |
| PLAU | -3,59853212 | 7,15128741 | 766,367155 | 1,6736E-23 | 6,0231E-20 |
| CDCP1 | -6,24586335 | 4,96049594 | 718,407152 | 3,4588E-23 | 8,0768E-20 |
| PKIA | -4,05544449 | 4,61024981 | 694,616312 | 8,1858E-23 | 1,4336E-19 |
| MECOM | -5,00728308 | 2,75719503 | 401,179029 | 1,2594E-22 | 1,9606E-19 |
| TMEM158 | -6,01271203 | 8,49160069 | 645,502732 | 2,1738E-22 | 3,0457E-19 |
| ARHGAP22 | -2,30600885 | 6,87142831 | 509,962336 | 7,0923E-21 | 9,0336E-18 |
| PRKG2 | -5,9647217 | 2,04457406 | 307,016047 | 2,2962E-20 | 2,298E-17 |
| TBX5-AS1 | -6,5161844 | 2,69255048 | 370,263718 | 2,7922E-20 | 2,6081E-17 |
| ANKRD29 | -3,14380408 | 3,8616196 | 401,780675 | 2,4079E-19 | 1,8743E-16 |
| PTGS1 | -4,32293506 | 5,73390258 | 388,130226 | 3,9468E-19 | 2,7649E-16 |
| COL10A1 | -5,42619213 | 2,74419591 | 261,924762 | 6,8395E-19 | 4,5632E-16 |
| SYT7 | -3,88673674 | 3,07269291 | 313,638855 | 9,8633E-19 | 6,2816E-16 |
| TRHDE-AS1 | -4,1387111 | 2,37603345 | 255,225572 | 1,9202E-18 | 1,1697E-15 |
| GNG11 | -1,66466833 | 8,61219227 | 343,566017 | 2,1983E-18 | 1,2833E-15 |
| IL13RA2 | -3,50298836 | 5,18084908 | 330,154244 | 3,9259E-18 | 2,0905E-15 |
| ESM1 | -4,51791683 | 5,13934285 | 329,885177 | 4,2213E-18 | 2,1123E-15 |
| RBPJ | -1,87164355 | 7,57130675 | 324,89157 | 4,8868E-18 | 2,361E-15 |
| DCBLD2 | -1,75645942 | 8,00459415 | 320,664101 | 5,888E-18 | 2,7499E-15 |
| ITGA2 | -3,31133532 | 7,24546569 | 315,636731 | 7,39E-18 | 3,3401E-15 |
| DUSP5 | -2,30522724 | 4,97442683 | 314,375952 | 7,8475E-18 | 3,436E-15 |
| HMGA2 | -2,10976875 | 6,90182036 | 309,236181 | 9,8704E-18 | 4,0675E-15 |
| CAMK2N1 | -2,42396795 | 7,76115882 | 306,487438 | 1,12E-17 | 4,3591E-15 |
| STEAP1 | -2,67366925 | 6,48178603 | 282,418312 | 3,5607E-17 | 1,2472E-14 |
| WNT16 | -4,06330443 | 5,27741099 | 273,478742 | 6,1194E-17 | 1,9486E-14 |
| NRIP3 | -2,238957 | 4,83804039 | 268,278837 | 7,3575E-17 | 2,2782E-14 |
| TRHDE | -4,08874404 | 3,53259233 | 255,46604 | 7,4797E-17 | 2,2782E-14 |
| GALNT6 | -1,82919544 | 7,17092606 | 265,138872 | 8,6288E-17 | 2,5187E-14 |
| ENSG00000290317 | -5,5960822 | 2,54807663 | 257,152761 | 9,544E-17 | 2,729E-14 |
| APCDD1 | -5,13317959 | 3,50126076 | 228,247439 | 1,2286E-16 | 3,4428E-14 |
| STC1 | -5,20013312 | 7,20040937 | 255,40019 | 1,4761E-16 | 3,9772E-14 |
| TAGLN3 | -5,93569192 | 2,1572465 | 212,805576 | 1,6108E-16 | 4,1794E-14 |
| SMIM3 | -1,94667927 | 6,58526409 | 252,422561 | 1,7138E-16 | 4,3657E-14 |
| SDCCAG8 | -1,30200016 | 6,1151062 | 251,453788 | 1,8078E-16 | 4,4681E-14 |
| BCL2A1 | -4,63536513 | 1,67238896 | 171,1997 | 1,8329E-16 | 4,4681E-14 |
| MMP1 | -4,24530527 | 10,3733915 | 251,031877 | 1,8496E-16 | 4,4681E-14 |
| DENND2A | -3,87264455 | 2,05372605 | 218,734376 | 2,3166E-16 | 5,5013E-14 |
| PHKA1 | -3,47764993 | 2,65681729 | 218,822352 | 3,0551E-16 | 7,0172E-14 |
| TGFA | -5,38836537 | 1,06472373 | 173,634107 | 3,6791E-16 | 8,0584E-14 |
| FZD8 | -1,74701531 | 5,60647103 | 234,789955 | 4,7053E-16 | 9,6951E-14 |
| WLS | -1,77886371 | 5,79819737 | 226,915306 | 7,4898E-16 | 1,478E-13 |
| ENSG00000289384 | -3,18526168 | 3,93615673 | 227,80593 | 8,2854E-16 | 1,6123E-13 |
| TBX5 | -5,79234593 | 1,7156223 | 183,321988 | 1,3602E-15 | 2,5076E-13 |
| NES | -3,0201768 | 5,3869921 | 215,568806 | 1,5222E-15 | 2,7698E-13 |
| FAM107B | -1,90023156 | 5,50210607 | 210,850724 | 2,0485E-15 | 3,5003E-13 |
| HECW1 | -3,47261075 | 1,88833118 | 163,00113 | 2,3187E-15 | 3,8675E-13 |
| BMAL2 | -1,90497719 | 6,05826365 | 207,135735 | 2,6064E-15 | 4,1974E-13 |
| PHLDA1 | -2,56103781 | 7,52033282 | 206,520487 | 2,7123E-15 | 4,3185E-13 |
| ADIRF | -2,25651275 | 10,4163093 | 205,288222 | 2,9421E-15 | 4,5299E-13 |
| BET1-AS1 | -2,96672477 | 4,11592536 | 204,099075 | 3,4309E-15 | 5,1992E-13 |
